# An essential protease, FtsH, influences daptomycin resistance acquisition in *Enterococcus faecalis*

**DOI:** 10.1101/2023.07.31.551240

**Authors:** Zeus Jaren Nair, Iris Hanxing Gao, Aslam Firras, Kelvin Kian Long Chong, Pei Yi Choo, Kevin Pethe, Kimberly A. Kline

**Affiliations:** Singapore-MIT Alliance for Research and Technology, Antimicrobial Drug Resistance Interdisciplinary Research Group, Singapore; Singapore Centre for Environmental Life Sciences Engineering, Nanyang Technological University, Singapore; Interdisciplinary Graduate Programme, Graduate College, Nanyang Technological University, Singapore; School of Biological Sciences, Nanyang Technological University, Singapore; Lee Kong Chian School of Medicine, Nanyang Technological University, Singapore; National Centre for Infectious Diseases (NCID), Singapore; Department of Microbiology and Molecular Medicine, University of Geneva, Geneva, Switzerland

**Keywords:** Daptomycin resistance, multiple peptide resistance factor (*mprF*), *ftsH*, *Enterococcus faecalis*, chaperones, *hrcA*

## Abstract

Daptomycin is a last-line antibiotic commonly used to treat vancomycin resistant Enterococci, but resistance evolves rapidly and further restricts already limited treatment options. While genetic determinants associated with clinical daptomycin resistance (DAP^R^) have been described, information on factors affecting the speed of DAP^R^ acquisition is limited. The multiple peptide resistance factor (MprF), a phosphatidylglycerol modifying enzyme involved in cationic antimicrobial resistance, is linked to DAP^R^ in pathogens such as methicillin-resistant *Staphylococcus aureus*. Since *Enterococcus faecalis* encodes two paralogs of *mprF* and clinical DAP^R^ mutations do not map to *mprF,* we hypothesized that functional redundancy between the paralogs prevents *mprF*-mediated resistance and masks other evolutionary pathways to DAP^R^. Here we performed *in vitro* evolution to DAP^R^ in *mprF* mutant background. We discovered that the absence of *mprF* results in slowed DAP^R^ evolution and is associated with inactivating mutations in *ftsH* resulting in the depletion of the chaperone repressor HrcA. We also report that *ftsH* is essential in the parental, but not in the Δ*mprF*, strain where FtsH depletion results in growth impairment in the parental strain, a phenotype associated with reduced glycolysis and reduced ability for metabolic reduction. This presents FtsH and HrcA as enticing targets for developing anti-resistance strategies.

**Graphical Abstract:** 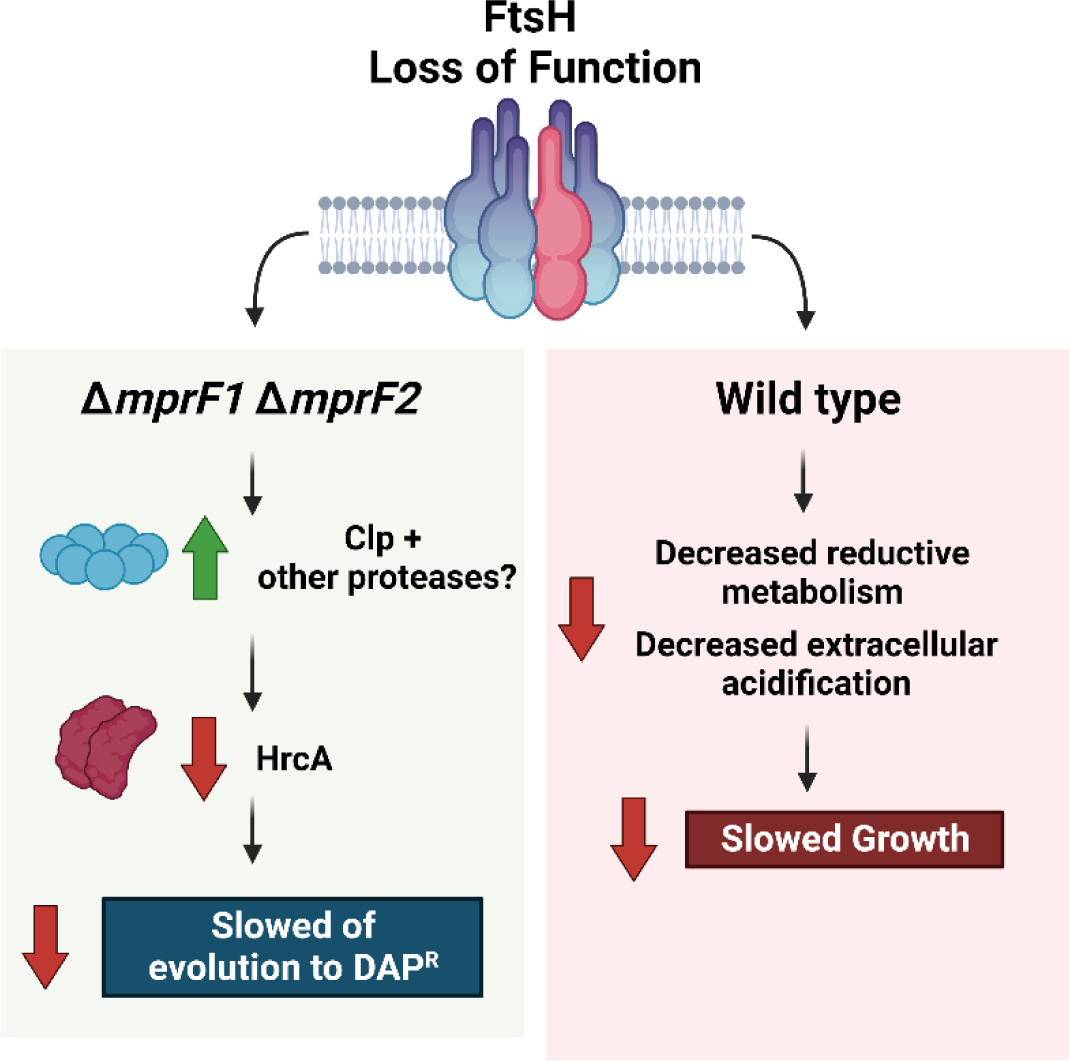

## Introduction

*Enterococci* are a major healthcare concern due to their association with hospital acquired infections (HAIs). Enterococci accounted for 14% of all HAIs in the USA from 2011-14 and 10% of HAIs in Europe in 2010, where *Enterococcus faecalis* comprise the majority of enterococcal HAIs, contributing up to 64.7% globally from 1997-2016 (Weiner et al., 2016, Zarb et al., 2012, Pfaller et al., 2019). Enterococci cause a variety of infections including catheter associated infections (CAUTI), endocarditis, peritonitis, colitis, diabetic foot ulcers, surgical site infections (Edmond et al., 1999, Hidron et al., 2008, Murdoch et al., 2009, Patterson et al., 1995, Weiner et al., 2016). Many of these infections are biofilm-associated, rendering them inherently more tolerant to antibiotics and difficult to treat (Ch’ng et al., 2019).

An additional challenge in treating Enterococcal infections is their intrinsic and acquired resistance to antimicrobials, including last line drugs such as vancomycin (Hollenbeck and Rice, 2012, Miller et al., 2014). For example, infections caused by vancomycin resistant Enterococci (VRE) are associated with increased mortality rates, lengthened hospital stays, and higher treatment and infection control costs (Reinseth et al., 2020, Miller et al., 2020, Carmeli et al., 2002, Prematunge et al., 2016, Mascini and Bonten, 2005, Song et al., 2003). Treatment of VRE infections typically involves the use of antibiotics of last resort such as linezolid and daptomycin (Patel and Gallagher, 2015). Daptomycin (DAP) is a lipopeptide antibiotic with broad activity against Gram-positive bacteria. It is positively charged when complexed with its calcium cofactor and targets the negatively charged bacterial membrane wherein it oligomerizes to cause membrane disruption, ion leakage, and eventual cell death (Steenbergen et al., 2005, Taylor and Palmer, 2016). Though DAP is typically effective in treating VRE infections, VRE can also acquire daptomycin resistance (DAP^R^), further reducing the already limited treatment options (Munita et al., 2014, Shoemaker et al., 2006, Arias and Murray, 2012, Kelesidis et al., 2011, Munoz-Price et al., 2005, Miller et al., 2020). While the rate of resistance of DAP in Enterococci is still relatively low at 0.1% for *E. faecalis* and 9% for *E. faecium*, DAP^R^ co-occurrence with VRE have been reported in several meta-analyses in Australia and New Zealand from 2007-18, where 15% of vancomycin resistant *E. faecium* are DAP^R^, and globally from 2003-10, 93.3% of Enterococcal DAP^R^ were VRE (Li et al., 2021, Kelesidis et al., 2011, Dadashi et al., 2021).

Given the clinical importance of DAP as a therapeutic and the emerging threat of resistance, enterococcal DAP^R^ associated mutations and resistance mechanisms have been characterized. Diverse genetic changes have been associated with DAP^R^ in both clinical isolates and *in vitro* settings. In vancomycin-resistant *E. faecalis* DAP^R^ patient isolates, mutations were identified in *liaF* (the negative regulator of the LiaFSR three-component system), *gdpD* (glycerophosphodiesterase) and *cls* (cardiolipin synthase), and similar mutations in all three genes were recapitulated in *in vitro* evolution of DAP-sensitive isolates to DAP^R^ (Palmer et al., 2011, Arias et al., 2011, Miller et al., 2013). Additionally, *in vitro* evolution also revealed DAP^R^-associated mutations not observed in clinical isolates such as in genes related to oxidative stress response (*gsh*, *yybT*, *selA*) and drug efflux (*mdpA*) (Miller et al., 2013). Similarly, mutations in *cls* and *liaFSR* have also been associated with DAP^R^ in *E. faecium* from both *in vivo* clinical isolates and *in vitro* evolution of DAP-sensitive strains to DAP^R^ (Tran et al., 2013a, Sinel et al., 2016, Wang et al., 2018, Li et al., 2022). Mutations in *liaF* as well as in *yvlB* (a putative LiaFSR target) in DAP^R^ strains suggest involvement of LiaFSR mediated membrane stress sensing (Arias et al., 2011, Miller et al., 2013). DAP^R^-associated mutations in *gdpD* and *cls*, decreased levels of phosphatidylglycerol (PG) and increased glycerophosphoryl-diglucosyldiacylglycerol (GPDGDAG), together with increased membrane rigidity and diversion of daptomycin away from the septum in DAP^R^ strains suggest that DAP^R^ is mediated through membrane remodeling (Arias et al., 2011, Mishra et al., 2012, Rashid et al., 2017, Tran et al., 2013b). Further investigation showed that LiaFSR is indeed one of the key systems that senses antimicrobials and initiates membrane remodeling to confer antibiotic resistance in *E. faecalis* (Khan et al., 2019). Taken together, these mutations suggest common DAP^R^ mechanisms among Enterococci involving antimicrobial stress sensing and membrane remodeling.

Despite the current advances in understanding DAP^R^ resistance mechanisms, information on factors that influence the rate of DAP^R^ acquisition is scarce. A complete understanding of DAP^R^ both in terms of factors directly affecting resistance, as well as factors that influence the speed and likelihood of progression towards resistance are equally important in the pursuit of anti-resistance strategies to mitigate potential widespread resistance in future.

DAP^R^ has been similarly well-studied in *Staphylococcus aureus*, where DAP^R^ is associated with *mprF* gain of function mutations and increased expression (Ernst et al., 2018, Mishra et al., 2009, Sabat et al., 2018, Sulaiman and Lam, 2021). Multiple peptide resistance factor (MprF) is a membrane bound enzyme that aminoacylates phosphatidylglycerol (PG) in the inner leaflet of the membrane and flips it to the outer leaflet, resulting in a reduction in the overall negative charge of the membrane and giving rise to electrostatic repulsion of cationic antimicrobials (Bao et al., 2012, Rashid et al., 2016, Ernst and Peschel, 2011). MprF mutations have not been reported in association with enterococcal DAP^R^. While there is only one *mprF* gene in the *S. aureus* genome, *E. faecalis* and *E. faecium* encode two paralogs– MprF1 and MprF2 – where MprF2 appears to be the major contributor to PG aminoacylation in *E. faecalis* (Bao et al., 2012, Rashid et al., 2023). We have also reported that *mprF* is closely tied to global lipidome regulation and cell physiology, and the absence of *mprF* significantly alters membrane lipid composition resulting in altered membrane fluidity, reduced secretion and increased dependence on exogenous fatty acids (Rashid et al., 2023). Given its daptomycin protective effects, we hypothesized that MprF redundancy afforded by its two encoding orthologues may mask additional daptomycin resistance events that occur during *in vitro* and *in vivo* evolution of *E. faecalis* to DAP^R^.

To investigate this possibility, we conducted *in vitro* evolution to DAP^R^ in *mprF* mutant backgrounds and discovered DAP^R^-associated mutations in several genes not previously associated with DAP^R^, including *ftsH*. FtsH is a conserved protease, and DAP^R^-associated mutations were only enriched in a Δ*mprF1* Δ*mprF2* background. Our data show that *ftsH* is essential in parental *E. faecalis* but not in the Δ*mprF1* Δ*mprF2* strain where its inactivation contributes to slowed evolution to DAP^R^. We found that FtsH indirectly affects levels of HrcA (the repressor of chaperone operons), which in turn influences the speed of DAP^R^ evolution. These findings provide evidence for a role of FtsH activity and HrcA in influencing DAP^R^ acquisition.

## Results

### Mutations in *ftsH* are enriched in Δ*mprF1* Δ*mprF2* during *in vitro* evolution to DAP^R^

Since MprF activity contributes to DAP^R^ in *S. aureus*, we investigated the contribution of the *E. faecalis mprF* paralogs to DAP^R^ (Ernst et al., 2018, Mishra et al., 2009, Sabat et al., 2018). While the *E. faecalis* OG1RF strain used in this study is DAP-sensitive (MIC of 4 – 8 µg/mL DAP), the loss of *mprF* makes it hypersensitive to DAP, reducing MIC by 2-4 fold in Δ*mprF2* and 4-fold in Δ*mprF1* Δ*mprF2* **(table 1)**. Hence, we hypothesized that the DAP-protective activity of MprF may mask resistance associated mutations not previously detected in DAP^R^ clinical isolates and *in vitro* evolution studies.

**Table 1.** Daptomycin Minimal Inhibitory Concentrations (MICs)

| **Table 1. Daptomycin Minimal Inhibitory Concentrations (MICs)** |  |
| --- | --- |
| **Strain** | **MIC in BHI (µg/mL)** |
| Wild type *E. faecalis* (OG1RF) | 4 – 8 |
| *ΔmprF1* | 4 – 8 |
| *ΔmprF2* | 2 – 4 |
| *ΔmprF1 ΔmprF2* | 2 |
| *ΔmprF1 ΔmprF2* passage control | 16 |
| *ΔmprF1 ΔmprF2 ftsH(G37X)* | 8 |
| *trePP::Tn* | 4 |
| *gelE::Tn* | 4 |
| *yckE::Tn* | 4 |
| *cryZ::Tn* | 4 |
| *hrcA::Tn* | 2 |
| *lutA::Tn* | 4 |
| *carB::Tn* | 4 |
| *dnaJ::Tn* | 4 |
| *groEL::Tn* | 8 |
| *ΔdnaK* | 4 |

To test this hypothesis, we performed *in vitro* evolution to DAP^R^ in *mprF* single and double mutants. Strains were first grown with DAP concentrations at 0.5X, 1X and 2X their respective MIC. The highest DAP concentration in which cultures grew was defined as the highest growth permissive concentration (HGPC) for the first round. In the following round, cultures were grown at 0.5X, 1X and 2X of the preceding day’s HGPC. This was done successively until an endpoint HGPC of 512 µg/mL DAP was achieved **(figure 1A)**. Using this approach, we were able to track the progression to DAP^R^ over time by recording the HGPC values. We observed that wild type and Δ*mprF1* reached the endpoint HGPC at similar rates and times, whereas Δ*mprF2* and Δ*mprF1* Δ*mprF2* progressed more slowly **(figure 1B)**. While Δ*mprF2* and Δ*mprF1* Δ*mprF2* reached endpoint resistance at similar times, Δ*mprF1* Δ*mprF2* displayed slower evolution in the initial phases from day 1 to 15 as compared to Δ*mprF2*. This can be explained in part due to the lower mutation rate measured for Δ*mprF1* Δ*mprF2* of 6.10 x 10^-9^ as compared to 3.80 x 10^-8^ for wild type, whereas Δ*mprF2* was more similar to wild type **(figure 1C)**.

**Figure 1.**
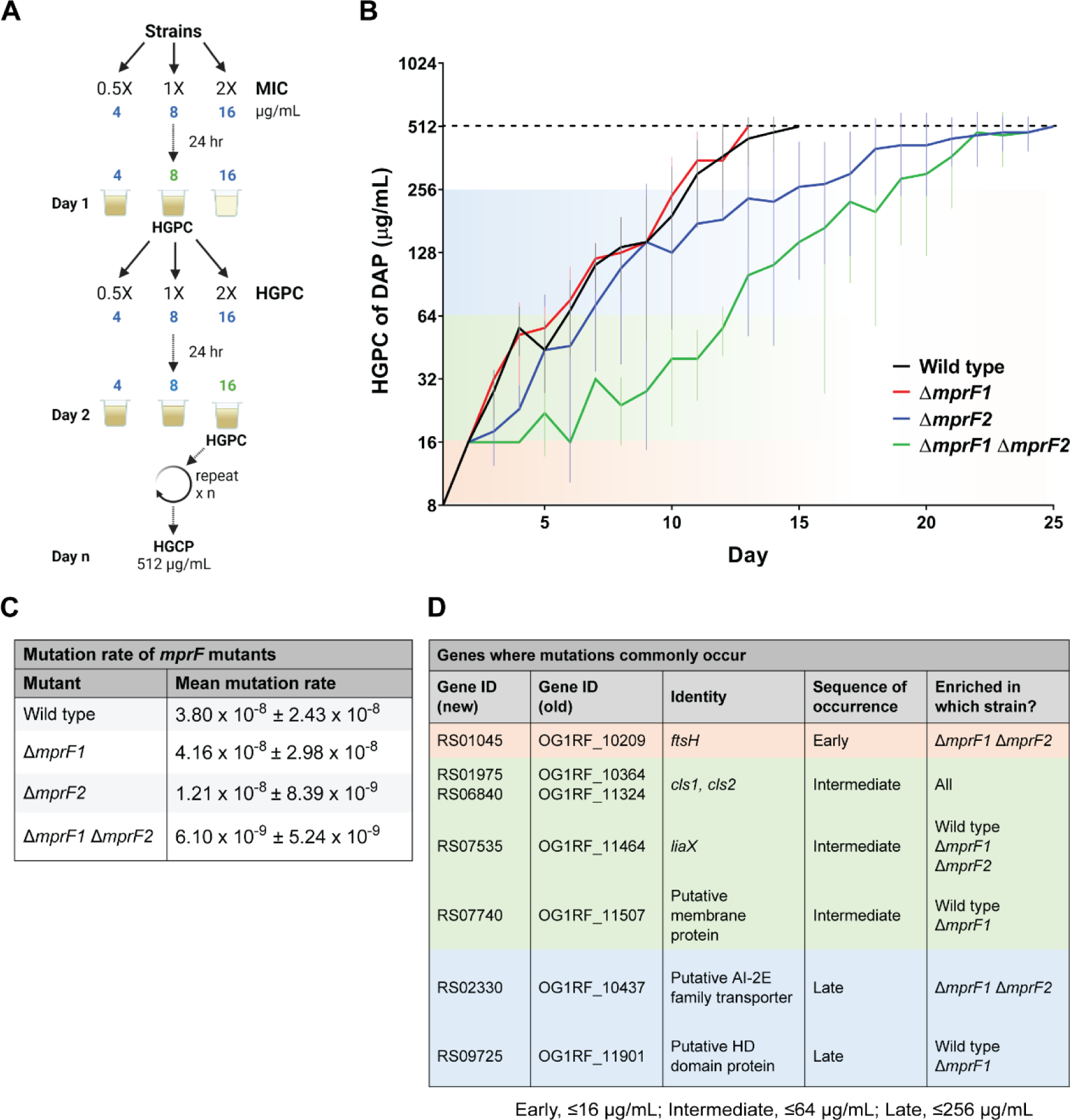
*In vitro* evolution of *mprF* mutants to DAP^R^ reveal novel mutations. **(A)** Workflow for *in vitro* evolution. Strains to be evolved were first grown in media supplemented with 0.5X, 1X or 2X their respective DAP MIC (e.g., 4, 8, 16 µg/mL) for 24 hours. The highest concentration of DAP that allowed for growth is termed the highest growth permissive concentration (HGPC) (e.g., 8 µg/mL). In the following passage, the HGPC culture was subcultured at 0.5X, 1X, 2X the HGPC of DAP (e.g., 4, 8, 16 µg/mL). This is repeated continually until an endpoint HGCP of 512 µg/mL of DAP was achieved. **(B)** The mean HGPC of daptomycin over time for each strain is plotted against time. Error bars indicate standard deviation from 8 parallel lines of evolution. **(C)** Mean mutation rate with standard deviation of *mprF* mutants assayed by the Luria-Delbrück fluctuation assay from 3 independent experiments. **(D)** Whole genome sequencing across evolution reveals an ordered progression of acquired mutations, with enrichment of some mutants in specific mutant backgrounds. The sequence of occurrence of mutations is based of DAP HGPC where the mutation first occurred (early, ≤ 16 μg/mL; intermediate, ≤ 64 μg/mL; Late, ≤ 256 μg/mL). Detailed information of all mutations observed are displayed relative to sequence of occurrence in evolution in **supplementary excel file S1A**, and relative to number of observed occurrences per gene in **supplementary excel file S1B**.

Clonal isolates were collected daily throughout evolution and sequenced to identify resistance-associated mutations in each genetic background **(figure 1D, supplementary excel file S1A, B)**. Mutations in cardiolipin synthase genes (*cls1*, *cls2*), previously implicated in *E. faecalis* DAP^R^ (Arias et al., 2011, Miller et al., 2013), emerged during the intermediate stages of evolution (DAP HGPC of 16-64 µg/mL) in all genetic backgrounds. We did not observe mutations in genes encoding the LiaFSR three-component system as described previously in DAP^R^ strains (Arias et al., 2011, Miller et al., 2013); however, mutations in the gene encoding LiaX – a antimicrobial sensing component for LiaFSR (Arias et al., 2011, Khan et al., 2019, Miller et al., 2013, Reyes et al., 2015) arose at a similar intermediate time in all genetic backgrounds except Δ*mprF1* Δ*mprF2* **(figure 1D)**. Mutations in *liaX* has recently been detected in DAP^R^ *E. faecalis* clinical isolates, but to our knowledge has yet to be detected in *in vitro* evolution screens (Ota et al., 2021). Mutations in genes encoding a predicted membrane protein (RS07740) and predicted HD domain protein (RS09725) were observed at intermediate and later stages of evolution (DAP HGPC > 64 µg/mL), respectively, and in the wild type and Δ*mprF1* background. Additionally, mutation in a predicted AI-2E family transporter (RS02330) was also observed at later stages of evolution (DAP HGPC ≤ 256 µg/mL) only in the Δ*mprF1* Δ*mprF2* background. Interestingly, mutations in *ftsH* were only observed in the Δ*mprF1* Δ*mprF2* background, during the earliest stages of evolution (DAP HGPC ≤ 16 µg/mL) **(figure 1D)**. FtsH is a conserved ATP-dependent zinc-metalloprotease and is membrane bound and hexameric in nature (Langklotz et al., 2012, Bieniossek et al., 2006).The cellular processes that FtsH influences are diverse across different organisms and depends largely on the substrates that it targets (Deuerling et al., 1997, Okuno and Ogura, 2013, Yepes et al., 2012, Kamal et al., 2019). FtsH was chosen for further investigation due to the intriguing phenomenon where mutations only occur in the Δ*mprF1* Δ*mprF2* background. The mutations in the other genes were not investigated in detail as their functions in DAP^R^ are either already known in the case of *cls*, and *liaX* or protein identity and detailed functions are not well defined in the case of the remaining genes RS07740, RS02330 and RS09725.

### FtsH is essential in the wild type but not Δ*mprF1* Δ*mprF2*

Many of the DAP^R^-associated *ftsH* mutations encode point mutations clustered within the ATP-binding site of the AAA+ domain or result in a G37X nonsense mutation near the N-terminal region of *ftsH*, suggesting that these mutations likely result in a loss of function **(figure 2A)**.To determine the contribution of *ftsH* mutations in DAP^R^ evolution, the *ftsH*(G37X) loss-of-function truncate was chosen for introduction into the wild type and Δ*mprF1* Δ*mprF2* backgrounds on a plasmid using a constitutive sortase A promoter (pGCP123-P*^srtA^*) **(figure 2A)**. Since FtsH forms homohexamers, we expected that FtsH(G37X) would assemble with native, chromosomally encoded FtsH, causing dominant negative dysfunction of the enzyme complex (Langklotz et al., 2012, Liu et al., 2022, Niwa et al., 2002). Indeed, expression of *ftsH*(G37X) resulted in slowed growth in wild type, but not Δ*mprF1* Δ*mprF2* mutant cells **(figure 2B)**. Within the wild type *ftsH*(G37X) expressing strain, log phase absorbance values were more variable than for the control strains **(figure 2B)**. We also noticed that small and large colony variants were only present in the wild type *ftsH*(G37X) expressing strain and subsequent analysis revealed loss of or reduced *ftsH*(G37X) insert sizes within the large colony variants but not the small colony variants **(figure 2C)**. These results suggest that FtsH loss of function (FtsH-LoF) is not tolerated in the wild type strain, but is permissible in Δ*mprF1* Δ*mprF2*. To confirm this, a proteolytically inactive *ftsH* variant – *ftsH*(H456Y) in which the conserved zincin motif within the protease active site was mutated as described by others – was constructed and placed under nisin inducible expression in a plasmid (pMSP3535-P*_nisA_*) (Arends et al., 2016, Bieniossek et al., 2006) **(figure 2A)**. As predicted, induction of 6his*-ftsH*(H456Y) in the wild type background resulted in slowed growth while expression of 6his-*ftsH* showed similar growth as the empty vector control. Moreover, attempts to introduce the *ftsH*(G37X) mutation into the chromosome were unsuccessful in the wild type background, but was possible in Δ*mprF1* Δ*mprF2* (data not shown). Taken together, these data show that *ftsH* is essential in a wild type background and its LoF is tolerated only in a Δ*mprF1* Δ*mprF2* genetic background.

**Figure 2.**
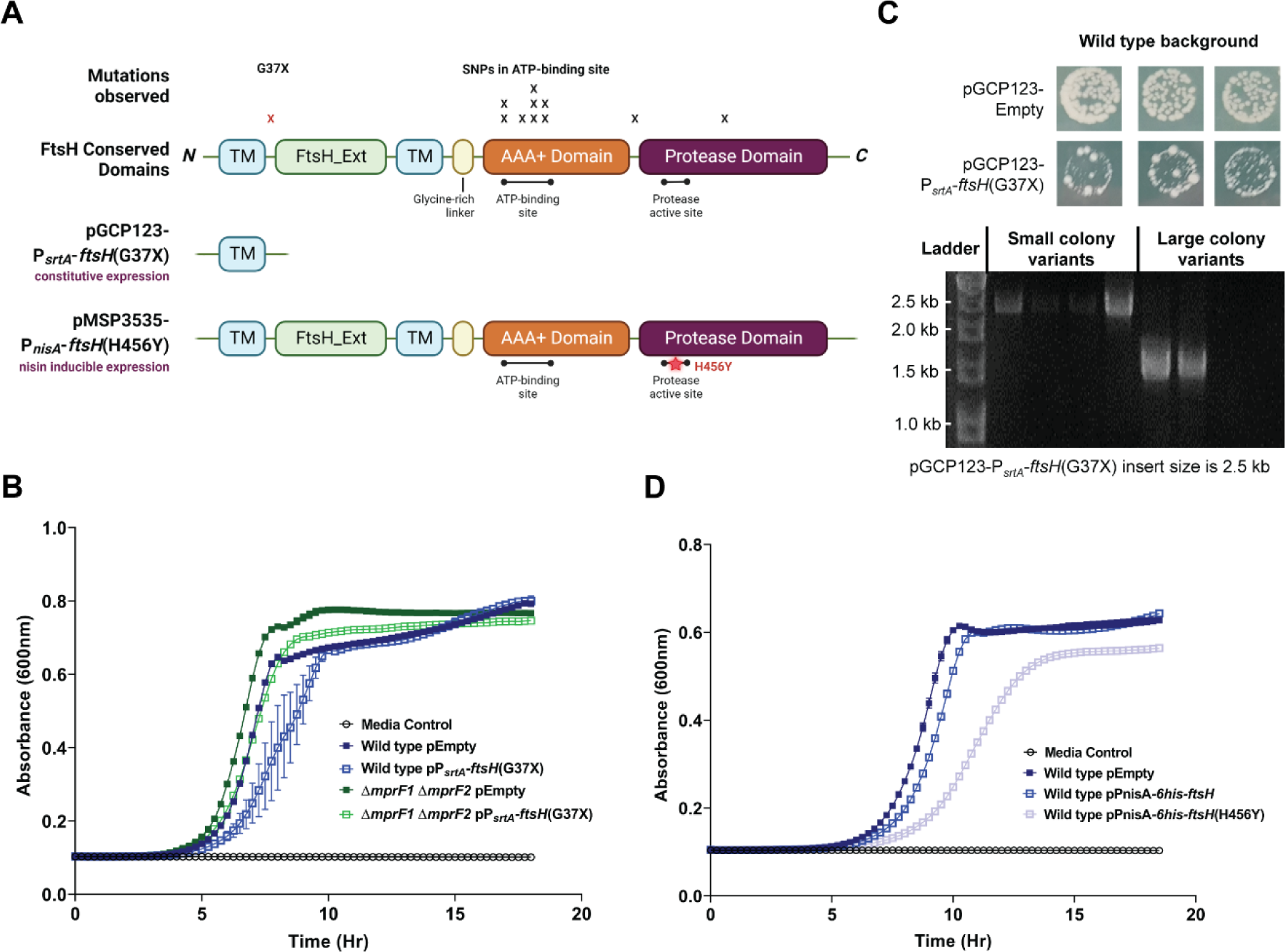
Mutations in *ftsH* are conditionally permissible in Δ*mprF1* Δ*mprF2*. **(A)** Map of single nucleotide polymorphism (SNP) mutations frequently observed in *ftsH* through evolution overlaid on where they occur with reference to conserved domains of FtsH. Diagrams showing the *ftsH* variants expressed – truncated FtsH(G37X) under constitutive expression (P*_srtA_*) or full length but proteolytically inactive FtsH(H456Y) under nisin inducible expression (P*^nisA^*). Created with BioRender.com. **(B)** Growth curves of wild type and Δ*mprF1* Δ*mprF2* with *ftsH*(G37X) expression. **(C)** *ftsH*(G37X) expression results in small and large colony variants in the wild type, where large colony variants show reduced/absent inserts within the expression vector. **(D)** Growth curves of the wild type with induced expression of either active *ftsH* or proteolytically inactive *ftsH*(H456Y).

### FtsH-LoF leads to metabolic impairments

To understand why an FtsH-LoF mutation caused a growth defect only in the wild type genetic background, we examined the viability of cells constitutively expressing *ftsH*(G37X) or inducibly expressing *ftsH*(H456Y). We observed similar proportions of propidium iodide (PI) stained cells in both populations, suggesting that membrane permeability is not affected by FtsH-LoF **(figure S1A)**. We also observed an increase in cell chaining upon FtsH-LoF in the wild type **(figure S1B)**.

To gain further insight into FtsH-dependent growth defects, we performed RNA sequencing following induced expression of either 6his-*ftsH* or 6his-*ftsH*(H456Y). However, we did not observe any obvious expression differences in genes that would explain this phenomenon **(supplementary excel file S1C)**. Despite the lack of insight from the transcriptomics data, we reasoned that the viability is unlikely to be affected since we observed no differences in PI staining and considered whether the slowed growth could be driven by altered metabolism. We therefore assessed the ability of the FtsH-LoF strains to reduce the resazurin dye to a fluorescent product using the Alamar blue assay as an indicator of electron flow in the membrane. We observed a decrease in fluorescence of the reduced resazurin dye in the wild type strain constitutively expressing *ftsH*(G37X) suggesting impairment in reductive metabolic activity **(figure 3A)**. This decrease was not observed following inducible expression of *ftsH*(H456Y), which could be due to differences in expression levels or due to added metabolic stress from the presence of nisin used for induction.

**Figure 3.**
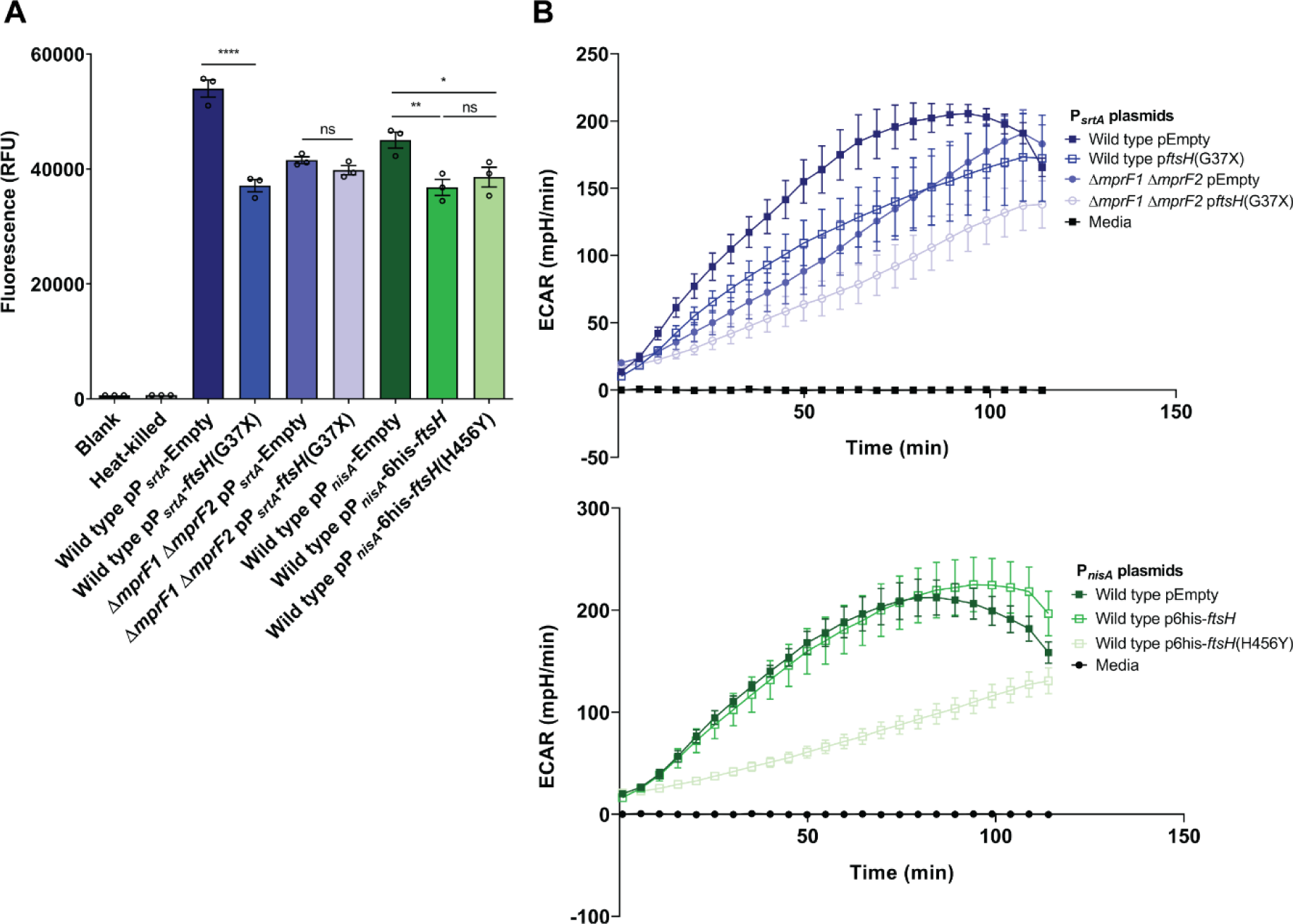
FtsH loss of function (FtsH-LoF) leads to metabolic changes. **(A)** Alamar blue assay measuring reductive metabolism of the FtsH-LoF strains. Larger relative fluorescence units (RFU) values indicate higher activity of metabolic reduction. Error bars represent the standard error of mean from 3 biological replicates. Tukey’s test for ANOVA. *, p<0.05; ****, p<0.0001. **(B)** Extracellular acidification rate (ECAR) quantified from *ftsH* loss of function strains using the Agilent Seahorse assay as an indirect measure of glycolysis. Error bars represent the standard error of mean from 4 biological replicates. Constructs in pP*_nisA_* plasmids are under a nisin inducible promoter induced with 25 ng/mL nisin.

We next considered the possibility that a shift in dominant cell metabolic pathways might explain the reduced metabolic activity. We performed Agilent Seahorse real-time cell metabolic analysis of extracellular acidification rates (ECAR) as an indirect measure of glycolysis, and oxygen consumption rate (OCR) as an indirect measure of oxidative phosphorylation in mid log phase cultures of wild type and Δ*mprF1* Δ*mprF2* expressing both constitutive and induced expression of inactive *ftsH* variants. FtsH-LoF correlated with reduced ECAR indicating reduced media acidification following expression of inactive *ftsH* variants **(figure 3B)**. This was observed for both constitutive and inducible expression of FtsH inactive variants, and in both the wild type and Δ*mprF1* Δ*mprF2*. Hence, these data indicate a generalized decrease in ability to acidify the media under FtsH-LoF **(figure 3B)**. The Seahorse OCR measurements were also largely similar across all strains indicating similar oxidative phosphorylation rates **(figure S2A)**. FtsH-LoF also did not result in any significant changes in quantified ATP levels **(figure S2B)**. Overall, these findings suggest that reduced growth of wild type cells expressing FtsH-LoF could be caused by reduced ability for metabolic reduction. Although the mechanism behind this phenomenon is unclear, we can rule out the contribution of oxidative phosphorylation and ATP production since they are similar across all strains.

### Speed of evolution to DAP^R^ is slowed under FtsH-LoF

Given that *ftsH* mutations were observed early in the slowed evolution of Δ*mprF1* Δ*mprF2* toward DAP^R^, we reasoned that these FtsH-LoF mutations might either be the reason for the slowed evolution or could be priming Δ*mprF1* Δ*mprF2* to acquire other DAP^R^ associated mutations in the later stages **(figure 1)**. Thus, the *ftsH*(G37X) mutation was introduced into the genome of Δ*mprF1* Δ*mprF2* for further investigation. As several days of passaging were carried out in low concentrations of DAP to encourage homologous recombination and retention of the *ftsH*(G37X) mutation in Δ*mprF1* Δ*mprF2*, a parallel culture of Δ*mprF1* Δ*mprF2* was passaged under the same conditions to serve as a control strain for comparisons in subsequent assays. This strain will be referred to henceforth as Δ*mprF1* Δ*mprF2* passage control. Δ*mprF1* Δ*mprF2 ftsH*(G37X) and Δ*mprF1* Δ*mprF2* passage control were subjected to *in vitro* evolution to DAP^R^ where we observed that Δ*mprF1* Δ*mprF2 ftsH*(G37X) evolved at a slower speed than Δ*mprF1* Δ*mprF2*, where under FtsH-LoF, the HGPC values were lower than the passage control at almost all time points and an additional 5 days was required to reach endpoint HGPC **(figure 4)**. This slower evolution is consistent with a one log lower mutation rate and 2-fold lower DAP MIC in Δ*mprF1* Δ*mprF2 ftsH*(G37X) as compared to Δ*mprF1* Δ*mprF2* passage control **(figure 4**, **table 1)**.

**Figure 4.**
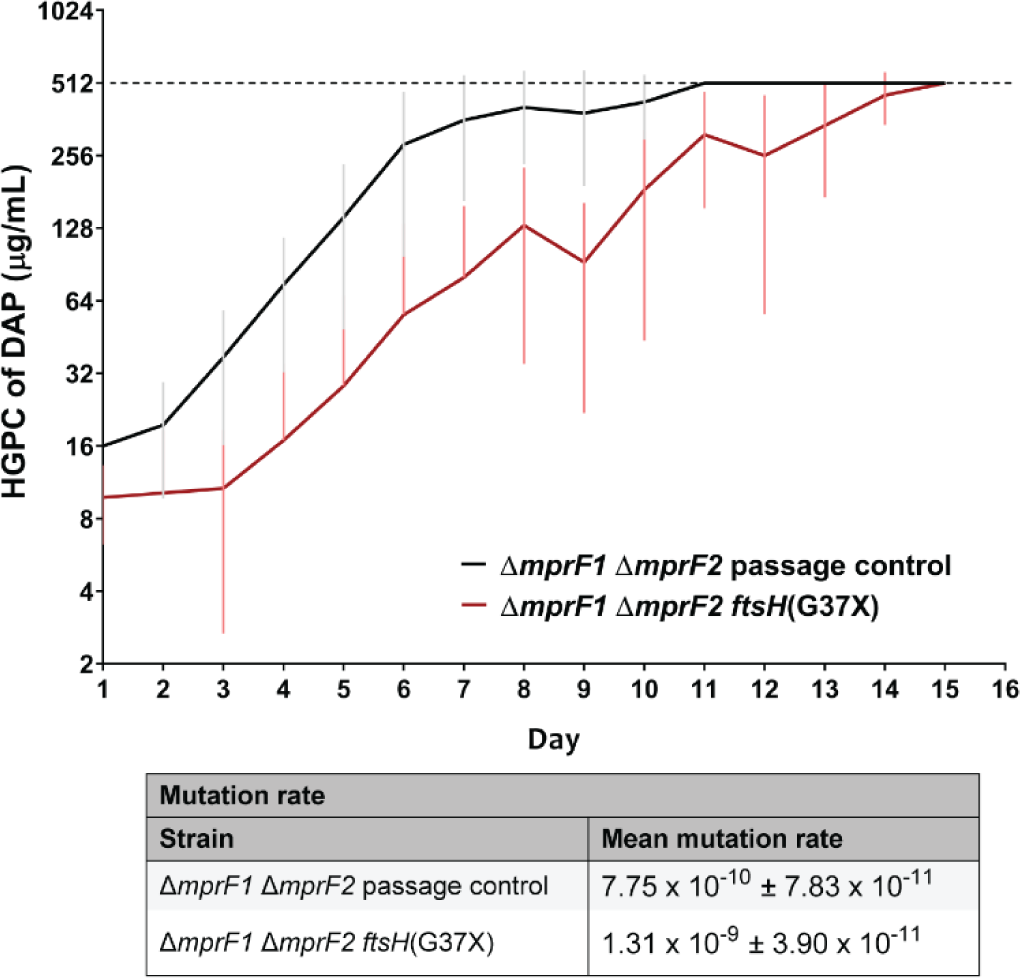
FtsH-LoF slows speed of evolution to DAP^R^ and decreases basal mutation rates in the Δ*mprF1* Δ*mprF2*. Mean highest growth permissive concentration (HGPC) of DAP across time from in vitro evolution of Δ*mprF1* Δ*mprF2 ftsH*(G37X) and Δ*mprF1* ΔmprF2 passage control to DAP^R^ of HGPC of 512 µg/mL DAP. Error bars indicate standard deviation from 9 parallel lines of evolution. Mean mutation rate assayed by the Luria-Delbrück fluctuation assay from 3 biological replicates where mean mutation rate is displayed together with standard deviation.

### Proteomic investigation of FtsH-LoF reveals *hrcA* as a key driver of slowed evolution

To determine the mechanisms underlying the slowed evolution and growth phenotypes following introduction of *ftsH*(G37X) to the *mprF* mutant background, we investigated the proteomic consequence of FtsH-LoF in the wild type background, by conducting peptide mass spectrometry on the whole cell lysates and membrane fractions of wild type pMSP3535-6his-*ftsH* and wild type pMSP3535-6his-*ftsH*(H456Y) following overnight induction with nisin. Proteomic changes common between the whole cell lysates and membrane were shortlisted, and proteomic changes that could be explained by transcriptomic differences were filtered out **(table 2)**. From this short list, we identified several proteins that were depleted following FtsH-LoF **(table 2)**, which may be explained by compensatory activity of other proteases such as the Clp protease which was transcriptionally induced when FtsH was non-functional (Log_2_FC for *clpP* = 1.63; *clpE* =1.34; *clpB* = 1.24, *clpC* = 1.15) **(supplementary excel file S1C).** Of the four proteins that accumulated in the FtsH-LoF strain **(table 2)**, ArcB and a putative amidase (RS02510) were verified to be substrates of FtsH by assessing protein stability and FtsH-dependent degradation under FtsH-LoF **(figure S3)**.

**Table 2.** Proteomic changes in FtsH-LoF (wild type p6his-FtsH(H456Y)) ^†^.

| Table 2. Proteomic changes in FtsH-LoF (wild type p6his-FtsH(H456Y))<sup>†</sup> |  |  |  |  |
| --- | --- | --- | --- | --- |
| Depleted Proteins |  |  |  |  |
| Gene | Gene Number (new; old ID) | Identity | Proteomic log<sub>2</sub>FC | RNAseq log<sub>2</sub>FC |
| *yckE* | RS05275; OG1RF_11014 | beta-glucosidase | -2.77 | - |
| *lysS* | RS01060; OG1RF_10212 | lysine-tRNA ligase | -2.69 | - |
| *pyrB* | RS07365; OG1RF_11430 | aspartate carbamoyltransferase | -2.54 | - |
| *lutA* | RS04635; OG1RF_10886 | putative lactate utilization Fe-S protein; homologous to *B. subtilis* *lutA* | -2.01 | 1.22 |
| *gelE* | RS07835; OG1RF_11526 | gelatinase E | -1.95 | - |
| *trePP* | RS12410; OG1RF_12425 | glycosyl hydrolase | -1.86 | - |
| *carB* | RS07350; OG1RF_11427 | carbamoyl-phosphate synthase large subunit | -1.69 | 0.67 |
| *cryZ* | RS07135; OG1RF_11383 | putative NADPH:quinone reductase | -1.67 | - |
| *hrcA* | RS05580; OG1RF_11076 | heat-inducible transcription repressor HrcA | -1.56 | - |
| Accumulated Proteins |  |  |  |  |
| Gene | Gene Number (new; old ID) | Identity | Proteomic log<sub>2</sub>FC | RNAseq log<sub>2</sub>FC |
| *arcB* | RS00500; OG1RF_10100 | ornithine carbamoyltransferase | 2.53 | -0.89 |
| RS08610 | RS08610; OG1RF_11679 | metal ABC transporter substrate-binding protein | 1.14 | -1.59 |
| *cls1* | RS01975; OG1RF_10364 | cardiolipin synthase 1 | 2.46 | - |
| RS02510 | RS02510; OG1RF_10473 | amidase | 2.36 | - |

<sup>†</sup> Proteomic changes that are not correlated to transcriptomic differences are displayed. Short list of proteomic changes which are common across membrane fractions and whole cell lysates. Refer to **supplementary excel file S1D, E** for full list of proteomic changes in membrane fractions and whole cell lysates respectively. Refer to **supplementary excel file S1C** for RNAseq data that was used to filter proteomic changes that were correlated with transcriptomic expression changes.

We hypothesized that the accumulation or depletion of these proteins might explain the slowed growth observed from FtsH-LoF in the wild type. We examined each of the depleted proteins either with transposon mutants (*yckE*::Tn*, lutA*::Tn*, gelE*::Tn*, trePP*::Tn*, carB*::Tn*, cryZ*::Tn*, hrcA*::Tn) (Kristich et al., 2008) or by CRISPRi silencing of genes for depleted proteins that were not available in the transposon library (*lysS* and *pyrB*) (Afonina et al., 2020). To mimic accumulation of proteins, *arcB,* RS08610*, cls1* and RS02510 were cloned into a nisin inducible plasmid for induced overexpression. This panel of mutants was assayed for growth and we observed that *trePP*::Tn, *lysS* silencing, and overexpression of RS02510 resulted in slowed growth, indicating that these gene products could be contributing to the growth defect observed in a wild type genetic background with FtsH-LoF **(figure S4A-F)**.

We next subjected the same panel of transposon mutants to *in vitro* evolution to DAP^R^ to determine their contribution to the slowed evolution in Δ*mprF1* Δ*mprF2 ftsH*(G37X). However, in the initial evolution assay all strains had a similar profile as wild type where the HGPC of every strain was saturated at the assay’s upper selection limit (2X HGPC) for most of the assay making it hard to distinguish any difference between the strains (data not shown). To overcome this limitation, evolution was performed at an expanded DAP selection range of 0.5X, 1X, 2X, 4X, 8X HGPC instead. We observed that only *hrcA*::Tn was significantly associated with slowed evolution **(figure 5A)**. However, when we calculated mutations rates, we found that *hcrA*::Tn was similar to wild type **(figure 5A)**. Of the remaining transposon mutants, most displayed similar evolution profiles as the wild type **(figure S5)**. The slight delay observed for *lutA*::Tn (lactate utilization protein) was due to a single outlier that evolved much slower than the rest **(figure S5A)**. Evolution of CRISPRi and overexpression mutants was not possible, due to plasmid insert loss during evolution despite maintenance of antibiotic selection pressure (data not shown). Hence, *hrcA* appears to play a major contributing role towards the slowed evolution in FtsH-LoF.

**Figure 5.**
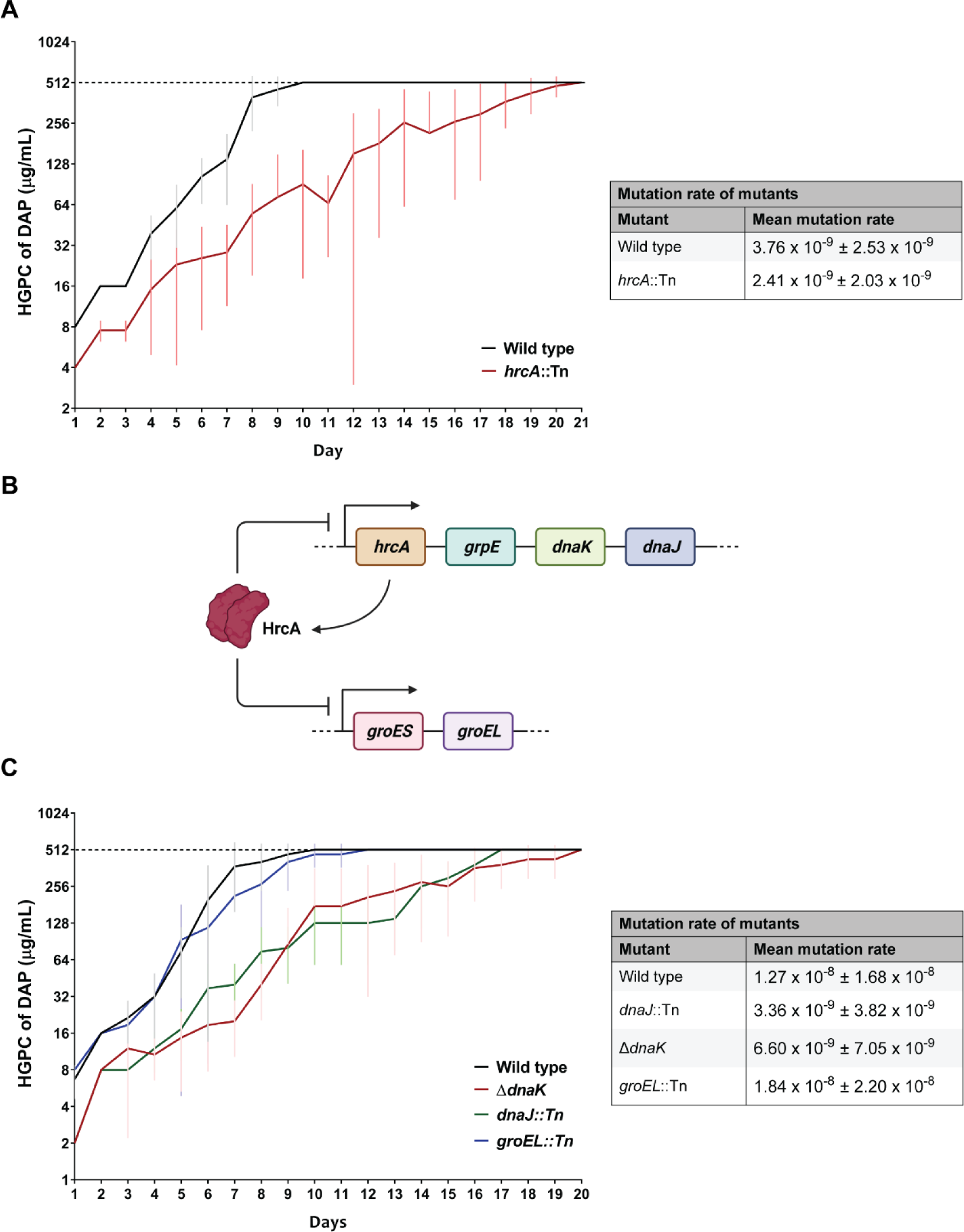
Disruption of HrcA and its target regulatory chaperones alters speed of DAP^R^ acquisition. Mean highest growth permissive concentration (HGPC) of DAP across time from *in vitro* evolution to DAP^R^ (HGPC of 512 µg/mL DAP) for **(A)** *hrcA*::Tn, and **(C)** chaperone mutants – Δ*dnaK*, *dnaJ*::Tn, *groEL*::Tn. Error bars indicate standard deviation from 9 parallel lines of evolution for **(A)** and 6 parallel lines of evolution for **(C)**. Evolution was performed using an expanded antibiotic selection range of 0.5X, 1X, 2X, 4X, 8X HGPC instead. Mean mutation rate on the right side of each graph assayed by the Luria-Delbrück fluctuation assay from 3 biological replicates where mean mutation rate is displayed together with standard deviation. **(B)** Model of the HrcA regulon, where HrcA negatively regulates the *hrcA-grpE-dnaK-dnaJ* and *groES-groEL* operons. Created with BioRender.com.

### Chaperones downstream of the *hrcA* regulon alter evolution speeds

HrcA is a transcriptional repressor of chaperone operons – *hrcA-grpE-dnaK-dnaJ* and *groES-groEL –* where it binds the conserved controlling inverted repeat of chaperone expression (CIRCE) element upstream of these operons (Schumann, 2016) **(figure 5B)**. From transcriptomic data of FtsH-LoF in the wild type background, we indeed observed upregulation of *grpE*, *dnaK* and *groEL* (Log_2_FC of 1.13, 1.53, 1.58 respectively) in concert with the depletion of HrcA **(supplementary excel file S1C)**. Hence, we reasoned that the depletion of HrcA could relieve transcriptional repression of these downstream chaperones, which could be contributing to the altered DAP^R^ evolution speeds. *dnaJ*::Tn, Δ*dnaK* and *groEL*::Tn were subjected to *in vitro* evolution using the same expanded DAP selection range as HGPC saturation at the upper limit of the assay occurred as described above (data not shown). We hypothesized that under FtsH-LoF, reduction of the repressor HrcA would result in upregulation of chaperones resulting in slowed evolution. Conversely, we expect that the loss of chaperone activity will result in a quickened evolution process.

However, unexpectedly, *groEL*::Tn displayed similar evolution profiles as the wild type strain, while evolution of *dnaJ*::Tn and Δ*dnaK* were slowed **(figure 5C)**. This slowed evolution could be due in part to the slightly lowered mutation rate and 2-fold reduction in DAP MIC of *dnaJ*::Tn and Δ*dnaK* **(figure 5C**, **table 1)**. Although we did not further investigate *grpE* and *groES*, we can expect evolution speeds to be similar to mutants of *dnaK* and *dnaJ,* and *groEL* respectively, since GrpE functions as a co-chaperone together with DnaK and DnaJ, while GroES and GroEL are co-chaperones that function together in the same complex (Harrison, 2003, Hayer-Hartl et al., 2016). While the *hcrA*-regulated chaperones that we tested displayed opposing phenotypes to what was expected, we speculate that their combined effects together with other HrcA-regulated genes result in the observed slowed evolution in loss of *hrcA*.

## Discussion

Treatment of Enterococcal infections has become increasingly challenging with the rise of antimicrobial resistance, including resistance to daptomycin which is one of the drugs of last resort used to treat drug resistant infections such as vancomycin resistant Enterococci (Patel and Gallagher, 2015). With *E. faecalis* contributing to the majority of Enterococcal infections, there is increasing interest to elucidate the factors driving DAP^R^ in this species (Pfaller et al., 2019). Through a combination of *in vitro* evolution assays and sequencing of DAP^R^ isolates, previous efforts have revealed membrane remodeling, antimicrobial stress sensing, oxidative stress response, and drug efflux to contribute to DAP^R^ (Arias et al., 2011, Miller et al., 2013, Khan et al., 2019, Mishra et al., 2012, Tran et al., 2013b, Tran et al., 2015).

However, less focus has been placed on factors affecting the speed of antibiotic resistance evolution, especially in the case of DAP^R^ where slowing resistance acquisition could inform anti-resistance strategies. Factors that broadly affect the propensity to evolve resistance to antibiotics have been well described. These factors influence antibiotic resistance evolution through DNA-repair machinery and stress response pathways, including the DNA-damage associated SOS-response, error-prone DNA polymerases, sigma factors, and the DNA translocase Mfd (Merrikh and Kohli, 2020, Ragheb et al., 2019, Al Mamun et al., 2012, Boshoff et al., 2003, Erill et al., 2007). Additionally, chaperones also provide buffering capacity for the fitness cost of resistance mutations affecting protein stability (Fay et al., 2021, Aguilar-Rodríguez et al., 2016). Other than these general factors, there are others that specifically influence evolution to DAP^R^. Recently, *liaFSR* was reported to affect the speed of DAP^R^ evolution in *E. faecium*, where deletion of *liaR* significantly slows evolution suggesting that LiaFSR activation is the dominant pathway to DAP^R^ in *E. faecium* (Prater et al., 2021). A synergistic effect of DAP with another antibiotic has also been reported to delay DAP^R^ acquisition where the co-administration of DAP with fosfomycin in *S. aureus* delayed the evolution to DAP^R^ (Mishra et al., 2022). While some information on factors affecting evolution to DAP^R^ exist, there is still limited mechanistic understanding at a genetic level for DAP^R^ acquisition in *Enterococci*.

Apart from the few well described mechanisms of DAP^R^ in *E. faecalis*, here we report that the multiple peptide resistance factor (MprF) also plays some role in DAP^R^ where the loss of *mprF2* hypersensitizes the already DAP-sensitive OG1RF strain by further decreasing DAP MIC. Through *in vitro* evolution of the *mprF* mutants to DAP^R^ to uncover novel mutations that might be otherwise masked by *mprF* activity, we discovered that evolution was slowed considerably in Δ*mprF1* Δ*mprF2*. Apart from this, by utilizing mutants of *mprF* we unmasked loss of function mutations (LoF) in *ftsH* observed only within the Δ*mprF1* Δ*mprF2* genetic background early in evolution **(figure 1D, 2A)**.

The effect of chaperones on accelerating protein evolution is well documented and is likely the reason for the observed slowed evolution in their absence. The DnaK chaperone can provide mutational robustness by buffering deleterious mutations that affect protein structure and function, and has been described to buffer the fitness cost of mutations associated with rifampicin resistance in *Mycobacteria* (Fay et al., 2021, Aguilar-Rodríguez et al., 2016). A similar mechanism might be at play in *E. faecalis* such that loss of *dnaK* leads to slowed resistance evolution due to reduced ability to buffer mutations that affect protein stability. DnaK has also been implicated in central metabolism and carbon source utilization in *E. coli* (Anglès et al., 2017). This could similarly be the case for *E. faecalis*, since we observed metabolic changes in terms of altered ability for extracellular acidification and metabolic reduction in the FtsH-LoF mutants where chaperones are upregulated **(figure 3, supplementary excel file S1C)**. DnaK and its co-chaperone DnaJ might also be essential in relieving *E. faecalis* of metabolic constraints that might be introduced by mutations accumulated through evolution.

Under FtsH-LoF, the resulting depletion of HrcA results in slowed evolution to DAP^R^. With HrcA being a chaperone operon repressor, it is expected that its depletion results in upregulation of downstream chaperones that contribute to this slowed evolution. Conversely, we would expect that disruption of these chaperones would enhance evolution instead. Unexpectedly, we instead observed slowed evolution when chaperones DnaK and DnaJ were disrupted. Hence at present, we do not yet fully understand how the depletion of HrcA enhances DAP^R^ evolution. One possibility is that by using the reductive approach in deleting or disrupting individual chaperones, we are only probing their individual contribution to resistance evolution which may not reflect the FtsH-LoF environment where multiple chaperones are upregulated under HrcA depletion. Furthermore, given that chaperones are canonically known to promote evolution by stabilizing deleterious mutations in proteins, it is unlikely that they are the sole reason behind the slowed evolution under HrcA depletion. It is more likely that the involvement of chaperones in combination with other *E. faecalis* genes regulated by HrcA together result in the slowed evolution under HrcA depletion, which is a topic for further investigation.

Additionally, HrcA depletion is unlikely to be the sole reason for the observed slowed evolution under FtsH-LoF since Δ*mprF1* Δ*mprF2 ftsH*(G37X) displayed lower mutation rates and this was not observed for *hrcA*::Tn. It is possible that the other accumulated proteins under FtsH-LoF might also play a role in slowing evolution, but we were unable to investigate further due to plasmid stability limitations in mimicking overexpression during *in vitro* evolution **(table 2)**. Additionally, with FtsH-LoF, there is a consequent decrease in HrcA, suggesting compensatory activation or upregulation of other proteases such as Clp that result in HrcA depletion. Another open question is the identity of these compensatory proteases that are responsible for HrcA depletion in FtsH-LoF. Nonetheless, our study provides evidence of the involvement of *ftsH* and *hrcA* in *E. faecalis* DAP^R^ evolution that has not been previously described and presents them as potential targets for means of influencing evolution rates and anti-resistance strategies. We also provide evidence of an alternative route to DAP^R^ involving protein quality control and chaperone regulation, apart from the well described routes involving antimicrobial stress sensing and membrane remodeling.

Our study also revealed that FtsH is essential in the wild type background but is dispensable in Δ*mprF1* Δ*mprF2*. Several proteins that were depleted or accumulated under FtsH-LoF could explain the reason for the slowed growth in the wild type, namely the depletion of TrePP, LysS and accumulation of a putative amidase (RS02510). Since TrePP is a glycosyl hydrolase responsible for the hydrolysis of glycosidic bonds, particularly that of trehalose-6-phosphate, and lysine-tRNA ligase is responsible for the ligation of lysine to tRNA, it is possible that reduced ability to break down complex sugars and produce essential lysine-tRNA could be contributing to the growth defect. Furthermore, the absence of MprF that utilizes lysine-tRNA as a substrate in Δ*mprF1* Δ*mprF2* could reduce the pressure of a limited lysine-tRNA pool allowing for normal growth under FtsH-LoF. Related to the growth defect, the accumulation of the putative amidase (RS02510) could also be contributing to the slowed growth, especially since amidases tend to play roles in cell division where they hydrolyze crosslinked peptidoglycan to allow for septation, dysregulation of this putative amidase could have similar effects (Vollmer et al., 2008, Do et al., 2020). Apart from growth related observations, FtsH-LoF in the wild type also resulted in a significant increase in cell chaining which could be mediated by the depletion of gelatinase E (GelE) under FtsH-LoF. Since gelatinases act to cleave autolysin to process it into its active form, the reduction in gelatinase E likely results in reduced autolysin activity resulting in dysfunctional cell division and increased cell chaining (Stinemetz et al., 2017). Furthermore, we have also shown that ArcB and RS02510 are substrates of FtsH, providing the first identification of FtsH substrates in *E. faecalis*. Therefore, while *ftsH* loss is tolerated in Δ*mprF1* Δ*mprF2*, it is essential in the wild type, where its loss leads to a growth defect driven by altered metabolism and altered cell division.

The reason behind the synthetic viability of FtsH-LoF in Δ*mprF1* Δ*mprF2* is still not fully understood. Given the altered lipidomic and metabolic landscape of Δ*mprF1* Δ*mprF2* (Rashid et al., 2023), it is possible that this would provide a permissive environment to offset the deleterious effects of FtsH-LoF. This could be mediated through glycosyl hydrolase (TrePP), where its disruption causes growth defects in the wild type and is depleted in FtsH-LoF. However, TrePP is transcriptionally up-regulated in Δ*mprF1* Δ*mprF2*, possibility compensating for the TrePP loss under FtsH-LoF (Rashid et al., 2023). While not known to affect growth, transcriptional and proteomic expression of *mprF2* was also increased in FtsH-LoF (Log_2_FC *mprF2* = 0.54, MprF2 = 2.59), which could be off-set by *mprF2* deletion in Δ*mprF1* Δ*mprF2*. While the picture is not yet complete, these findings hint towards an altered metabolic environment within Δ*mprF1* Δ*mprF2*, which is an avenue for future investigation.

Taken together, we have demonstrated that FtsH is essential in wild type *E. faecalis*, but loss of function is permissible in Δ*mprF1* Δ*mprF2* which slows DAP^R^ evolution through the indirect depletion of HrcA and subsequent changes in regulatory flux of the downstream chaperone operons **(figure 6)**. Under FtsH-LoF in the wild type background, the ability for metabolic reduction and extracellular acidification is reduced along with FtsH-LoF associated changes in TrePP, LysS and amidase levels resulting in growth impairment. Whereas in FtsH-LoF in Δ*mprF1* Δ*mprF2*, growth is not affected, instead, HrcA is indirectly depleted by compensatory action of other proteases. While the loss of HrcA results in slowed evolution, the contribution of downstream genes in the regulon, *dnaK* and *dnaJ* does not fully explain the cause. It is likely that there is more complex higher order regulation present involving other genes that results in a net decrease in evolution speeds. This study provides the first major characterization of FtsH both in terms of its substrates and its functional role in *E. faecalis* involving growth and metabolism, as well as the previously undescribed involvement in antibiotic resistance and resistance acquisition. The possibility of manipulating DAP^R^ evolution by targeting FtsH, HrcA and chaperone presents an enticing opportunity for their utility as both a research tool and as possible candidates for development of anti-resistance strategies. This is especially the case for FtsH where its essentiality further highlights its potential as a therapeutic target.

**Figure 6.**
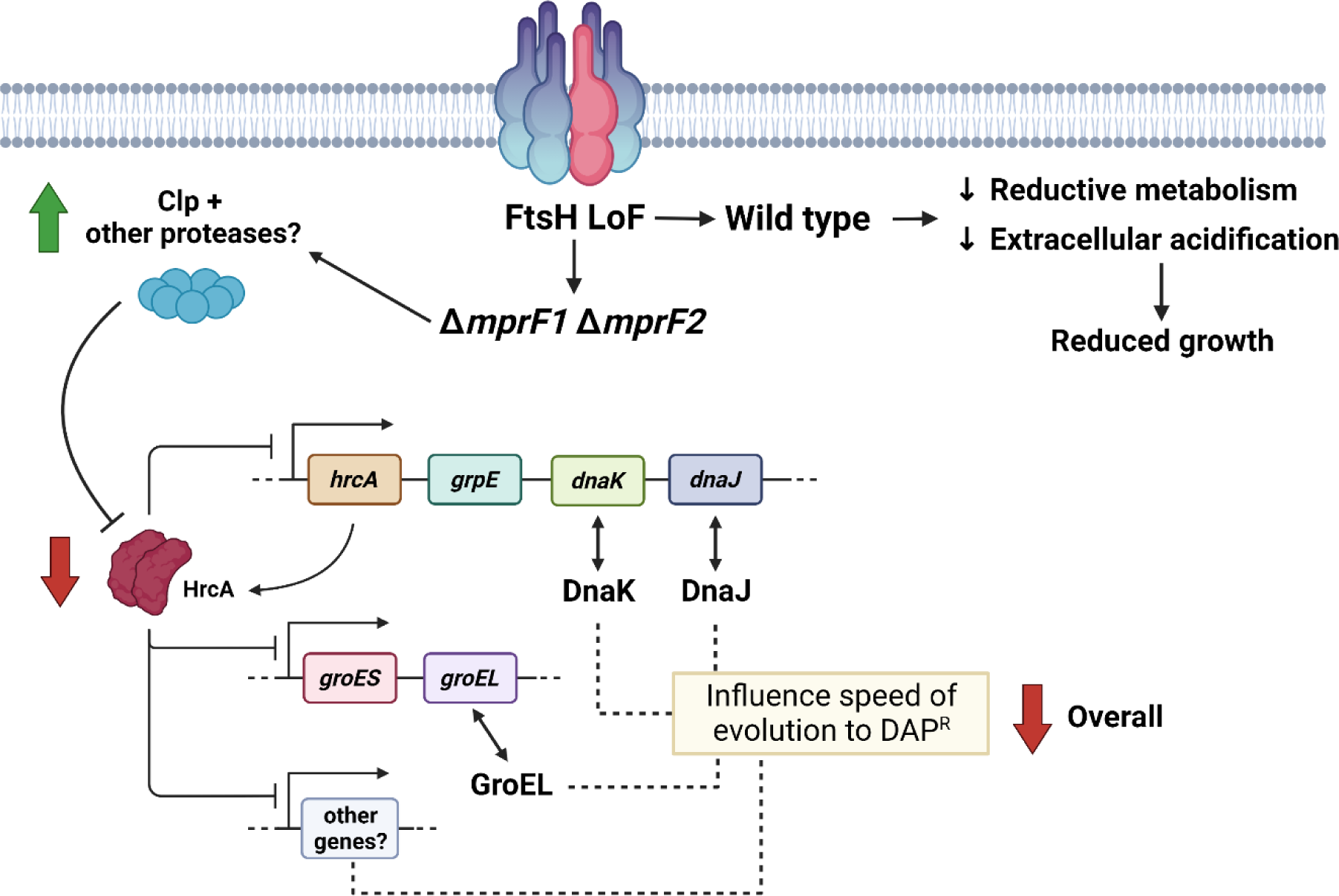
Model of FtsH’s influence on speed of DAP^R^ evolution and reduced growth in the wildtype. FtsH loss of function (LoF) indirectly leads to an increase in the Clp protease. This along with other proteases likely results in depletion of HrcA which relieves repression of the chaperone operons (*hrcA-grpE-dnaK-dnaJ* and *groES-groEL*). Further investigation revealed that *dnaK*, *dnaJ* influence the speed of DAP^R^ evolution. This combined with the extended regulatory effects of *hrcA* on other genes likely results in an overall combined effect of decreased evolution speed. FtsH-LoF also results in metabolic changes such as decreased ability for acidification and reductive metabolism that could be partly responsible for the reduced growth in the wild type. Created with BioRender.com.

## Experimental procedures

Strains, growth conditions, growth kinetics, live/dead staining, RNA sequencing and cloning methods are detailed in **supplementary text**.

### *In vitro* evolution of *E. faecalis* to daptomycin resistance

The protocol was adapted from a previously published *in vitro* evolution experiment done in *E. faecalis* V583 (Palmer et al., 2011). For each strain, multiple parallel lines of evolution experiment were performed. First, 100X dilutions of overnight bacterial cultures of each strain were made in BHI supplemented with 1.25 mM calcium chloride (Sigma, USA) (50 mg L^-1^ Ca^2+^) containing daptomycin (DAP) (Gold Biotechnology, USA) concentrations of 1X MIC, 2X MIC and 4X MIC, and incubated at 37 °C in static conditions for 22 to 26 hrs. Cultures of every evolution line were examined for visible bacterial growth. Bacterial cultures at the highest growth-permissive concentrations (HGPCs) are diluted 100X into fresh DAP-containing medium at 0.5X, 1X and 2X HGPC. This was repeated until HGPC of 512 μg mL^-1^ was achieved. Bacterial cultures were then passaged in plain BHI broth for 3 days to obtain stable mutants. Isolates were glycerol stocked each day in 25 % v/v glycerol. Refer to **supplementary figure S6** for the schematic of the *in vitro* evolution workflow. In instances where evolution profiles of the tested strains are consistently at the 2X HGCP and are saturated at the upper HGPC limit of the assay, an expanded range of 0.5X, 1X, 2X, 4X and 8X HGCP is used for selection instead.

### Whole genome sequencing

Whole genome sequencing was conducted on the glycerol stocked isolates from the wild type, Δ*mprF1*, Δ*mprF2* and Δ*mprF1* Δ*mprF2* backgrounds. Genomic DNA was extracted from overnight bacterial cultures using PureLink Genomic DNA Mini Kit (Thermo Fisher Scientific). Library preparation using MiSeq v3 and whole genome sequencing using MiSeq was done by the sequencing facility of Singapore Centre of Life Science Engineering (SCELSE, Singapore). Data was analyzed using CLC Genomics Workbench 8.0. The complete OG1RF reference genome (NC_017316) from NCBI database was used for mapping and annotation. The threshold variant frequency was set as 35 %. Non-synonymous mutations within coding regions were filtered for. All structural variations were manually confirmed on the mapping track.

### Minimal inhibitory concentration (MIC) by microplate dilution

Stationary phase cultures to be tested were grown until mid-log phase and normalized to OD_600_ of 0.7. MIC assays were performed in a 96-well plate as described previously (Wiegand et al., 2008), with the following modifications. Antibiotics were diluted in BHI media supplemented with 1.25 mM calcium chloride (50 mg L^-1^ Ca^2+^), in 2-fold dilutions, from 256.0 µg mL^-1^ to 0.5 µg mL^-1^ of daptomycin. Cultures with daptomycin were incubated for 16-18 hrs at 37 °C in static conditions before assessing for growth in the wells to estimate the MIC.

### RNA sequencing

Sequencing of RNA was done from wild type pMSP3535-6his-*ftsH*(H456Y) and wild type pMSP3535-*ftsH*(H456Y). Detailed methods are described in the **supplementary text** file.

### FtsH proteomic analysis

Wild type pMSP3535-6his-*ftsH*(H456Y) and wild type pMSP3535-*ftsH*(H456Y) strains were grown to mid-log phase and induced for expression of their respective plasmids’ gene constructs with 125 ng mL^-1^ of nisin for 16-18 hrs at 37 °C in static conditions and cell pellets were harvested. The membrane fraction was isolated from the harvested pellets as previously described, resuspended with 100 μL of 50 mM Tris-HCl, pH 8.0, and boiled with 33.3 μL of NuPAGE^®^ LDS Sample Buffer (4X) (Thermofisher, USA) and 10 μL of 1 M DTT (Maddalo et al., 2011). Samples were then run on SDS-PAGE on a 4-12 % NuPAGE^®^ Bis-Tris mini gel in a XCell SureLock^®^ Mini-Cell filled with MES SDS running buffer (Invitrogen, USA) until samples just entered the gel. Gels were then silver-stained by fixing with 50 % v/v methanol and 5 % v/v acetic acid solution, sensitizing with 0.02 % w/v sodium thiosulfate solution, silver-stained with 0.1 % w/v silver nitrate and 3 % v/v formalin solution and developed using 2 % w/v sodium carbonate and 1.5 % v/v formalin solution. The concentrated protein band of each lane was excised and stored in Eppendorf tubes filled with water. Samples were then sent to the Taplin Mass Spectrometry Facility, Harvard Medical School, Boston, Massachusetts, USA for peptide mass spectrometry and proteomic analysis. Peptide counts were normalized using tweeDEseq (TMM normalization) and statistics were done using Reproducibility-Optimized Test Statistic (ROTS) (Esnaola et al., 2013, Suomi et al., 2017).

### Mutation rate assay (Luria-Delbrück fluctuation assay)

Overnight stationary phase cultures were diluted 10, 000X in 40 mL of BHI supplemented with 1.25 mM calcium chloride (50 mg L^-1^ Ca^2+^). 100 μL of diluted culture was then added into each well of a 96-well microtiter plate, sealed and incubated at 37 °C in static conditions for 16-18 hrs. 24 wells from the plate were pooled followed by serial dilution and plating on non-selective BHI agar plate for CFU enumeration. This determines the average cell number (N). Whole volumes (100 μL) of each of the 72 wells/cultures were then transferred into wells of a 24-well microtiter plate containing 900 μL BHI supplemented with 1.25 mM calcium chloride and daptomycin (dilution was taken into account such that final daptomycin concentration is 3X MIC). Plates were incubated at 37 °C in static conditions and observed for growth visually by the presence of turbid wells for up to 7 days. The fraction of wells/cultures with zero growth indicating zero mutant cells is defined as *p_0_*. The expected number of mutation events per culture (m) is calculated as, *m* = −ln (*p*_0_). The mutation rate (μ) is calculated as: 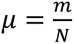.

### Alamar blue assay

Overnight cultures were normalized to OD_600_ 0.5 in PBS and diluted 1:10. The ability to reduce the resazurin dye was measured using the AlamarBlue™ HS cell viability reagent (Thermoscientific, USA) according to the manufacturer’s instructions.

### Seahorse assay

Mid-log phase cultures were washed once and normalized to OD_600_ 0.7 in BHI. Cultures were then added to a Cell-Tak^TM^ Cell and Tissue Adhesive (Corning, USA) coated XF96 cell culture microplate (Agilent, USA). Sterile media was added as blanks for background measurement. Microplate wells were coated with 25 µL of 22.4 µg mL^-1^ Cell-Tak^TM^ prior to use according to the manufacturer’s instructions. Plates were then centrifuged at 6, 000 x g for 15 mins to allow for cells to adhere to the bottom of the plate. A XFe96 sensor cartridge that has been soaked in calibration solution according to the manufacturer’s instruction was first loaded into the Seahorse XFe96 Analyzer (Agilent, USA) for instrument calibration, followed by the microplate containing the adhered cultures. Cultures were then measured for their oxygen consumption rate (OCR) and extracellular acidification rate (ECAR) for 120 mins at the following cycle settings: Mix – 2 mins 30 s, Wait – 0 mins, Measure – 4 mins, no injection.

## Data availability

Whole genome sequence files are available on NCBI, Sequence Read Archive (SRA) (Accession: PRJNA830756). RNA sequencing files are available on NCBI, Gene Expression Omnibus (GEO) (Accession: GSE201323) and SRA (Accession: PRJNA830869).

## Acknowledgements

We thank Cristina Colomer-Winter for critical reading of the manuscript and Swaine Chen for important scientific input over the course of this project. We would like to thank Ekaterina Sviriaeva from Lee Kong Chian School of Medicine, Nanyang Technological University for access, training, and the initial consumables for the Seahorse XFe96 Analyzer. We would also like to thank Ross Tomaino from Taplin Mass Spectrometry Facility, Harvard Medical School, Boston, Massachusetts for providing support for peptide mass spectrometry. We also extend our appreciation to Gary Dunny from University of Minnesota Medical School for providing the *E. faecalis* OG1RF transposon library, for which several mutants were used in our assays. We also thank Kline lab member Qingyan Chen, for her assistance with constructing wild type pMSP3535-P*_nisA_*-*cls1*.

This work was supported by the National Research Foundation and Ministry of Education Singapore under its Research Centre of Excellence Programme, as well as by the Singapore Ministry of Education under its Tier 1 program (MOE2017-T1-001-269) and the National Medical Research Council Open Fund (MOH-000645), both awarded to K.A.K and transferred to K.P.

## Author contributions

Conceptualization: ZJN, IHG, KAK

Formal analysis: ZJN, IHG, AF, KKLC

Funding acquisition: KAK

Investigation: ZJN, IHG, AF, KKLC, PYC

Methodology: ZJN, IHG, AF, KKLC, PYC

Project administration: ZJN, KAK

Supervision: ZJN, KP, KAK

Writing – original draft: ZJN, KAK

Writing – review & editing: ZJN, IHG, AF, KKLC, PYC, KP, KAK

## Abbreviated Summary

FtsH, a conserved protease, influences daptomycin resistance evolution rates in *Enterococcus faecalis* by indirectly affecting the availability of the chaperone repressor HrcA. FtsH is also essential in the wild type genetic background where its loss of function results in altered metabolism in terms of decreased extracellular acidification and ability for metabolic reduction resulting in slowed growth. However, FtsH loss is well tolerated in the multiple peptide resistance factor (MprF) mutant Δ*mprF1* Δ*mprF2*.

