## Supplementary figures and tables for "An essential protease, FtsH, influences daptomycin resistance acquisition in *Enterococcus faecalis*"

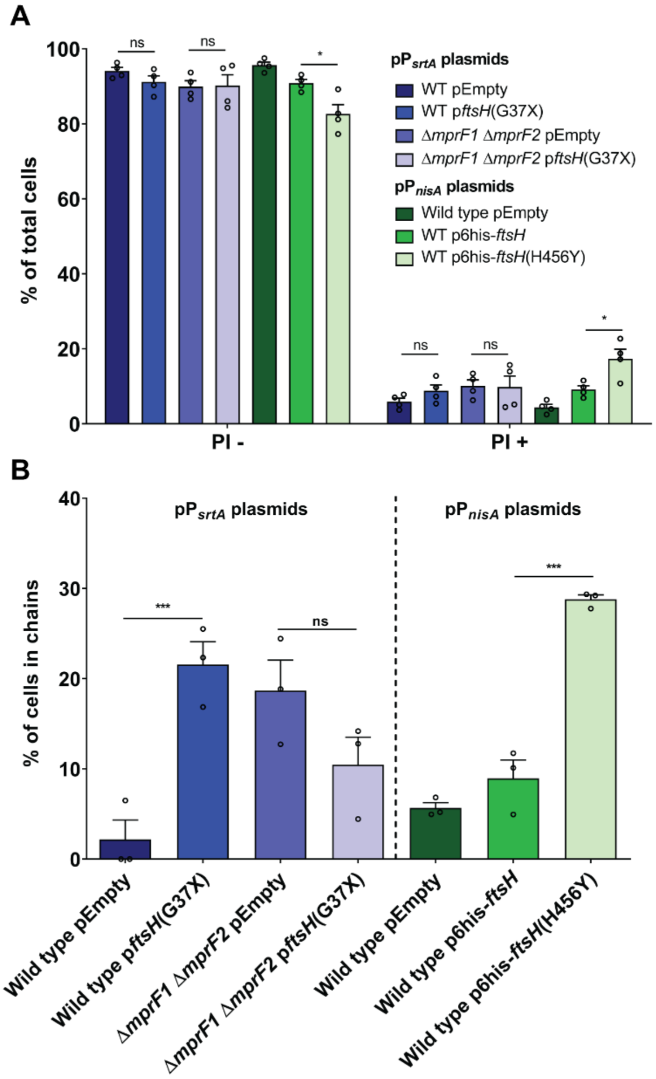


**Figure S1. FtsH loss of function (FtsH-LoF) has a minor effect on cell viability and increases cell chaining in the wild type background. (A)** Live-dead staining on stationary phase cultures with proportion of propidium iodide (PI) stained cells displayed. Error bars represent the standard error of mean from 4 biological replicates. At least 100 cells per replicate were analyzed. Tukey's test for ANOVA. *, $p<0.05$. **(B)** Enumeration of cells in chains from phase contrast microscopy. Chaining cells are defined as 3 or more cells adjacent to each other. Error bars represent the standard error of mean from 3 biological replicates. At least 100 cells per replicate were analyzed. Tukey's test for ANOVA. ***, $p<0.001$. Constructs in pP$_{nisA}$ plasmids are under a nisin inducible promoter induced with 25 ng/mL nisin.

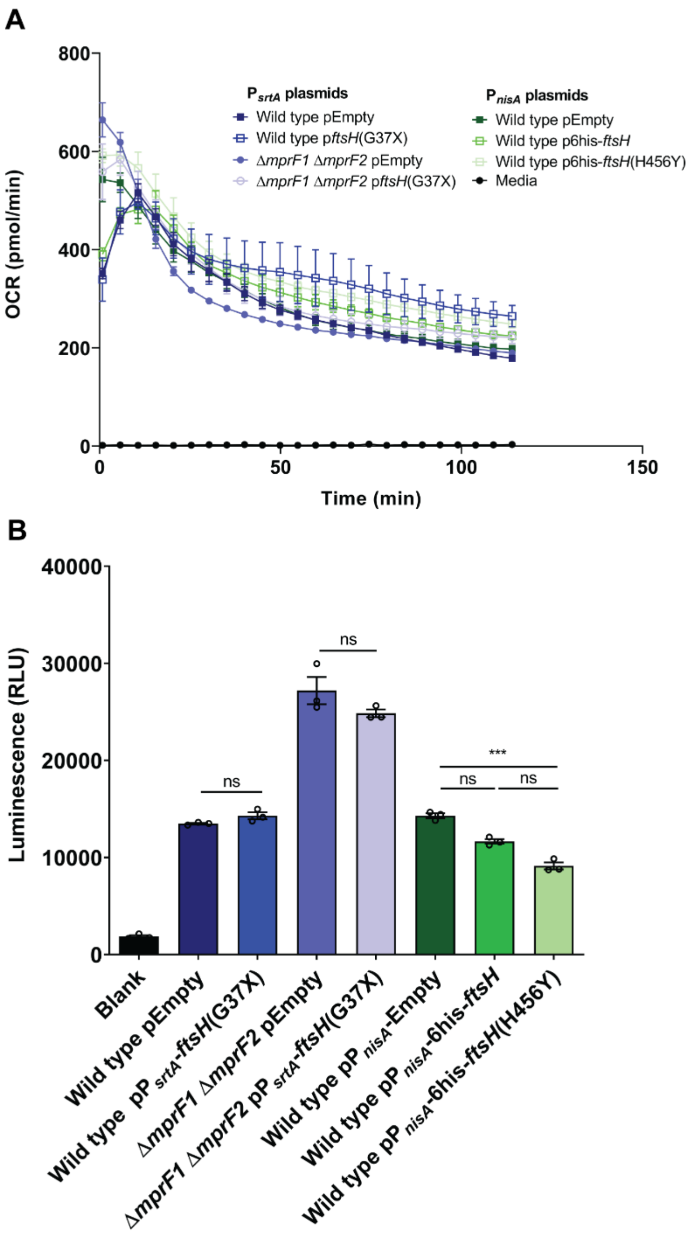


**Figure S2**. **FtsH-LoF does not affect oxygen consumption rate or ATP production. (A)** Oxygen consumption rate (OCR) quantified from *ftsH* loss of function strains using the Agilent Seahorse assay as an indirect measure of oxidative phosphorylation. Error bars represent the standard error of mean from 4 biological replicates. Constructs in pP$_{nisA}$ plasmids are under a nisin inducible promoter induced with 25 ng/mL nisin. **(B)** Luciferase-based ATP quantification of the FtsH loss of function strains. Luminescence measured is reflective of the amount of ATP present. Error bars represent the standard error of mean from 3 biological replicates. Tukey's test for ANOVA. ***, p<0.001.

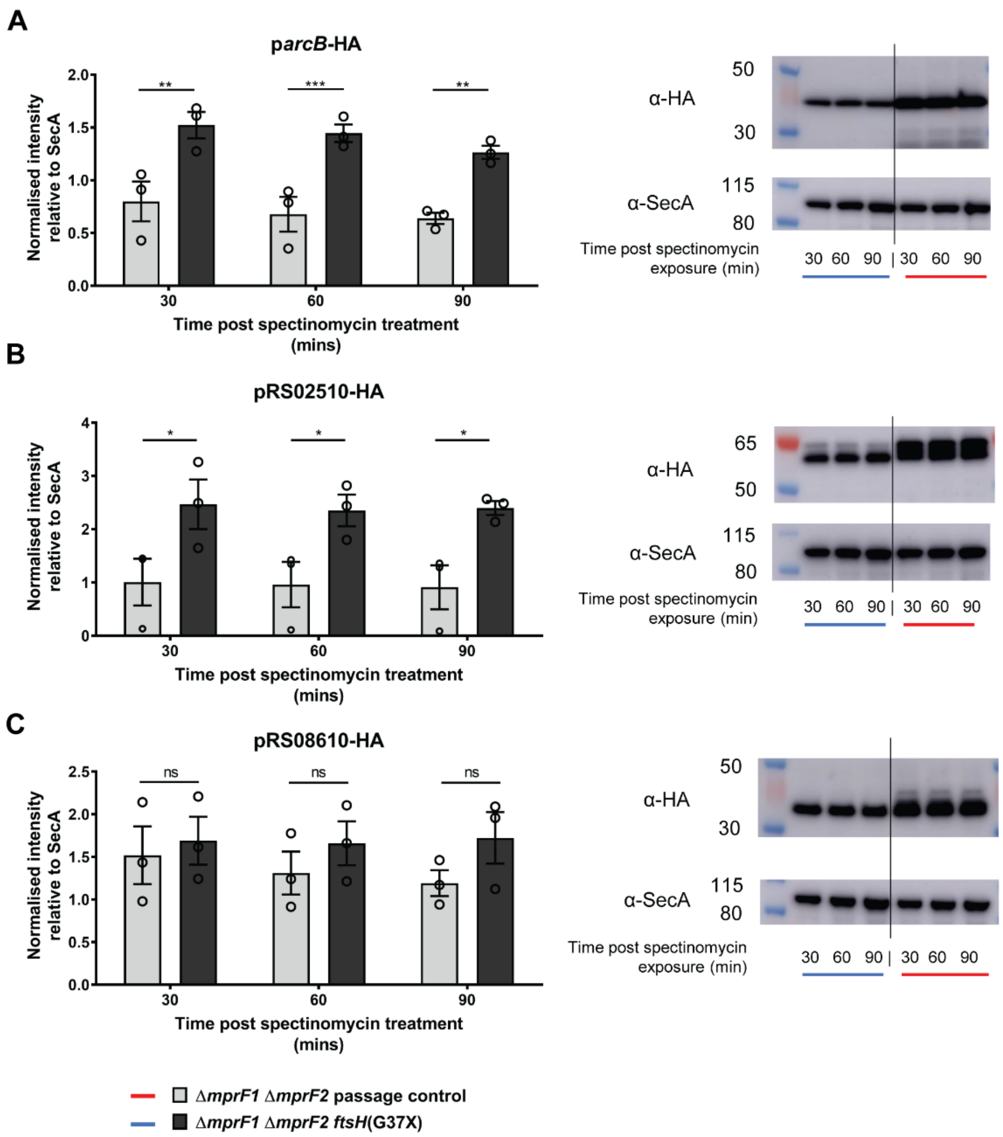


**Figure S3. ArcB and RS02510 (amidase) are likely substrates of FtsH.** Putative substrates – **(A)** ArcB, **(B)** RS02510, **(C)** RS08610 - were tagged with HA and harbored in the pMSP3535-P= nisin inducible expression vector and assayed for FtsH-dependent degradation using the Δ*mprF1* Δ*mprF2* passage control and Δ*mprF1* Δ*mprF2* strains. 125 ng/mL nisin was used for induction. Strains were treated with spectinomycin to block protein synthesis and protein abundance was monitored by western blot across 90 minutes post treatment. Blots for SecA were used as loading controls. Estimated protein sizes: ArcB-HA = 38.6 kDa, RS02510-HA = 59.6 kDa, RS08610-HA = 35.2 kDa, SecA = 96.2 kDa. Densitometry for protein abundance is displayed on the left of the respective blots. Error bars represent the standard error of mean from 3 biological replicates. Fisher's LSD test for two-way ANOVA. *, $p<0.05$; **, $p<0.01$; ***, $p<0.001$.

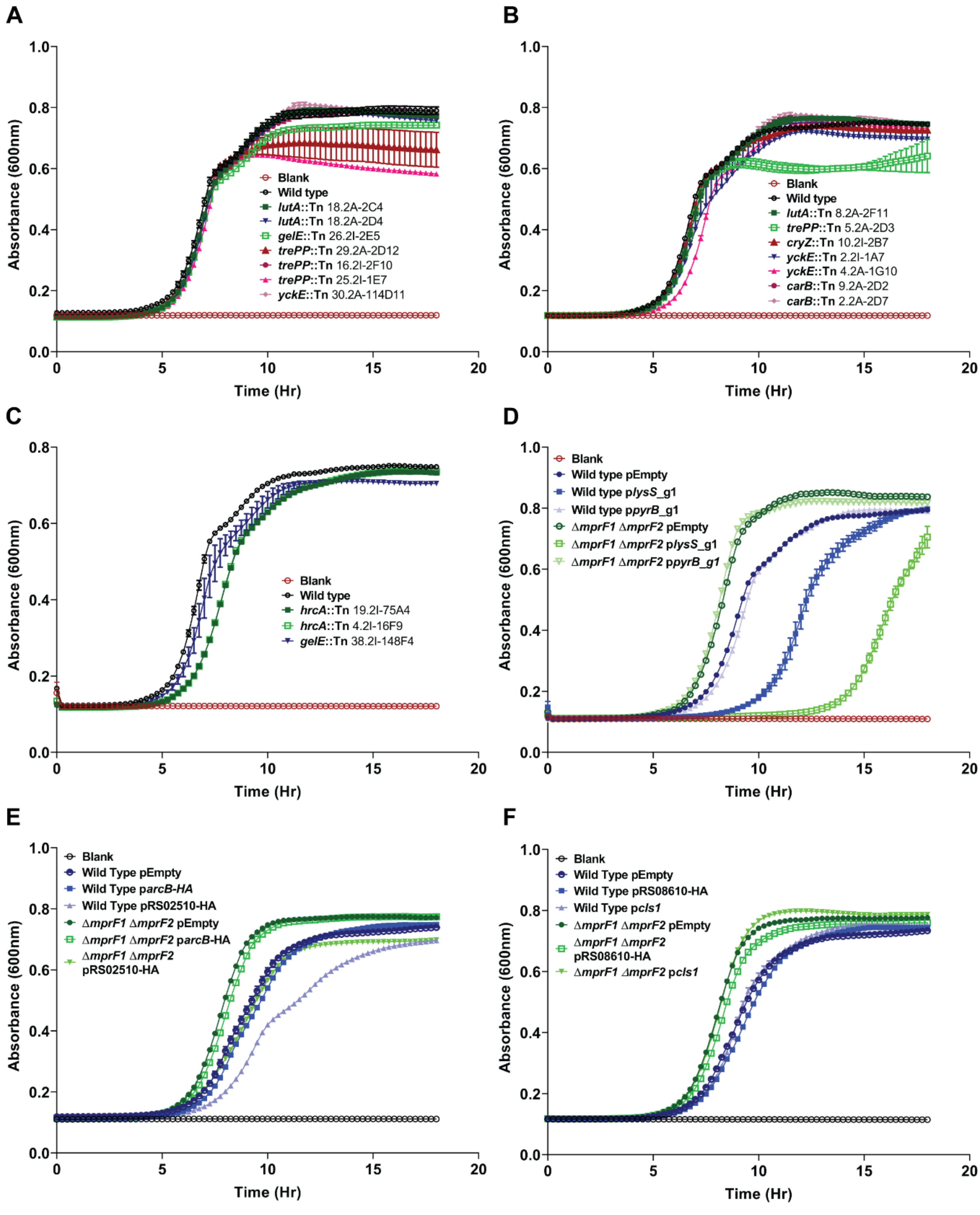


**Figure S4. Growth curves of mutant strains of depleted and accumulated proteins from proteomic analysis as well as wild type treated with varying concentrations of chaperone inhibitors. (A, B, C)** Transposon (Tn) mutants of depleted protein tested assayed individually for growth. **(D)** For depleted proteins where no Tn mutants are available, CRISPRi knockdowns of the respective genes were done instead, and growth assayed. **(E, F)** For accumulated protein hits, overexpressing strains of the respective genes were used, and growth assayed. Genes to be overexpressed are placed under a nisin inducible plasmid pMSP3535 and induced with 200 ng nisin for overexpression.

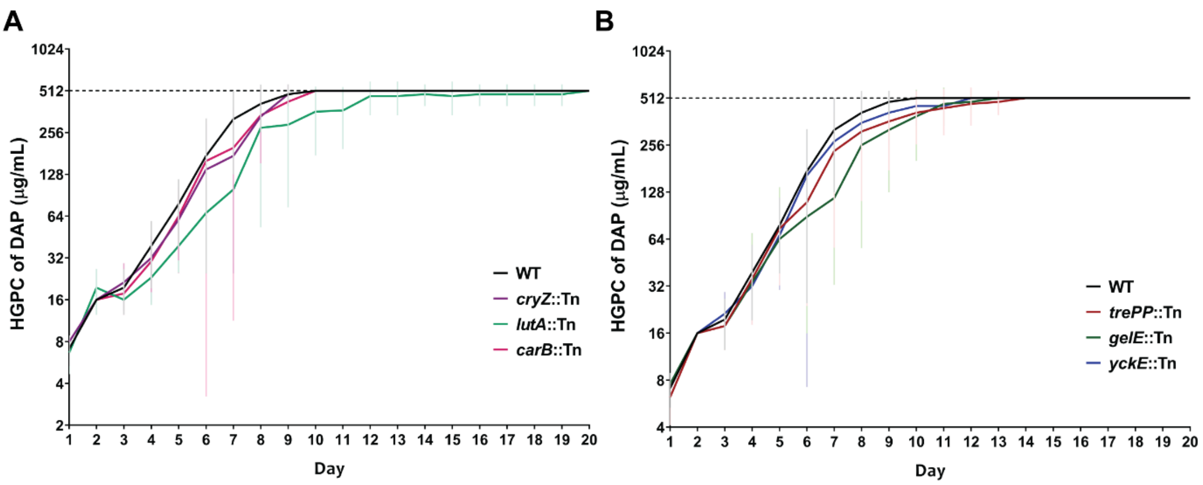


**Figure S5. *In vitro* evolution to DAP$^{R}$ in Tn mutants corresponding to other depleted proteins in FtsH-LoF in the WT.** Highest growth permissive concentration (HGPC) of DAP across time from *in vitro* evolution to DAP$^{R}$ (HGPC of 512 µg/mL DAP) for **(A)** *cryZ*::Tn, *lutA*::Tn, *carB*::Tn, and **(C)** *trePP*::Tn*, gelE*::Tn*, yckE*::Tn. Error bars indicate the standard deviation from 9 parallel lines of evolution. Evolution was performed using an expanded antibiotic selection range of 0.5X, 1X, 2X, 4X, 8X HGPC instead.

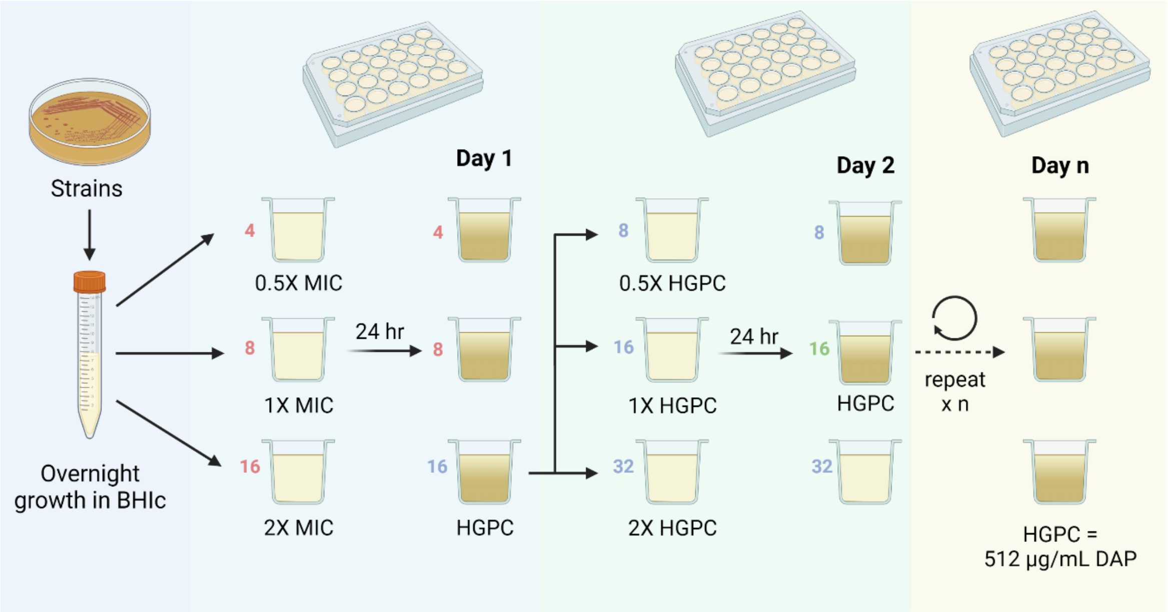


**Figure S6. Schematic showing the process of *in vitro* evolution.** Strains to be tested are first grown overnight in BHI supplemented with calcium (BHIc) to create the starter culture. The starter culture is then inoculated into wells of a 24-well plate containing media with daptomycin (DAP) that is 0.5X, 1X and 2X MIC of that respective culture. Following overnight growth, the highest concentration of DAP in which the culture was able to grow is taken as the highest growth permissive concentration (HGPC). This culture is used as the inoculum for the next subculture where it is grown at 0.5X, 1X and 2X HGPC of DAP. This process continues until a final HGPC of 512 µg $mL^{-1}$ of DAP is reached. Created with BioRender.com.

The colored numbers adjacent to the wells are example values to illustrate how a typical *in vitro* evolution might progress. [i.e., Cultures grown in 4, 8 and 16 µg $mL^{-1}$ of DAP in the first day where after 24 hours of growth, day 1's HGPC is 16 µg $mL^{-1}$. In the second day, cultures from HGPC of day 1 was subcultured into 8, 16, 32 µg $mL^{-1}$ of DAP where after 24 hours of growth, day 2's HGPC is 16 µg $mL^{-1}$. This carries on for n number of days until HGPC of 512 µg $mL^{-1}$ of DAP is reached].

### Supplementary tables

| Table S1. Bacteria Strains and Culture Conditions | | | |
|---|---|---|---|
| **Bacterial Strains** | **Relevant Information / Genotype** | **References / Source** | **Media Used** |
| | ***E. coli*** | | |
| Stellar™ Competent Cells (*E. coli* HST08) | *F–, endA1, supE44, thi-1, recA1, relA1, gyrA96, phoA, Φ80d lacZΔM15, Δ(lacZYA - argF) U169, Δ(mrr - hsdRMS - mcrBC), ΔmcrA, λ–* | Clontech, Takara Bio Inc., Japan | - |
| DH5α pGCP123-$P_{srtA}$ | DH5α harbouring pCYW2*; $Kan^r$<br>*pCYW2: pGCP123 parent plasmid (Nielsen et al., 2012) with PsrtA promoter. Expression vector. | Lab stock | LB, Miller broth (BD, USA) with 50 µg/mL kanamycin (Thermo Scientific, USA) |
| DH5α pMSP3535-$P_{nisA}$ | DH5α harbouring pMSP3535*; $Erm^r$<br>*pMSP3535: Shuttle vector for nisin-controlled inducible expression. | pMSP3535 was a gift from Gary Dunny, University of Minnesota (Addgene plasmid # 46886) (Bryan et al., 2000) | LB, Miller broth with 300 µg/mL erythromycin (Sigma Aldrich, USA) |
| Stellar™ Competent Cells pMSP3545-d*cas9* | *E. coli* HST08 harbouring pMSP3545-d*cas9**; $Erm^r$<br>*pMSP3535-dcas9: Shuttle vector for nisin-inducible expression of dcas9 | (Afonina et al., 2020) | LB, Miller broth with 300 µg/mL erythromycin |
| DH5α pABG5.2mini*<br>(*alternative name: pGCP123) | DH5α harbouring pABG5.2mini*<br>*pABG5.2mini: Shuttle expression vector | (Nielsen et al., 2012) | LB, Miller broth with 50 µg/mL kanamycin |
| TOP10 pGCP213 | TOP10 harbouring pGCP213*; $Erm^r$<br>*pGCP213: Temp-sensitive shuttle vector used for allelic replacement in E. faecalis | (Nielsen et al., 2012) | LB, Miller broth with 500 µg/mL erythromycin |

| | | | |
|---|---|---|---|
| Stellar™ Competent Cells pGCP213-*ftsH*(G37X) | *E. coli* HST08 harbouring pGCP213-*ftsH*(G37X); $Erm^r$ | This study | LB, Miller broth with 500 µg/mL erythromycin |
| Stellar™ Competent Cells pGCP213-Δ*dnaK* | *E. coli* HST08 harbouring pGCP213- Δ*dnaK*; $Erm^r$ | This study | |
| ***E. faecalis*** | | | |
| OG1RF | $Fus^r$, $Rif^r$, wild-type strain | American Type Culture Collection (ATCC© 47077™) | BHI (Neogen, USA) |
| OG1RF Δ*mprF1* | $Fus^r$, $Rif^r$, Δ*mprF1* | (Kandaswamy et al., 2013) | |
| OG1RF Δ*mprF2* | $Fus^r$, $Rif^r$, Δ*mprF2* | (Kandaswamy et al., 2013) | |
| OG1RF Δ*mprF1* Δ*mprF2* | $Fus^r$, $Rif^r$, Δ*mprF1*Δ*mprF2* | (Rashid et al., 2023) | |
| OG1RF Δ*mprF1* Δ*mprF2* *ftsH*(G37X) | $Fus^r$, $Rif^r$, Δ*mprF1*Δ*mprF2, ftsH*(G37X) | This study | |
| OG1RF Δ*mprF1* Δ*mprF2* DAP-passage control | $Fus^r$, $Rif^r$, Δ*mprF1*Δ*mprF2,* passage control strain | This study | |
| OG1RF Δ*dnaK* | $Fus^r$, $Rif^r$, Δ*dnaK,* passage control strain | This study | |
| OG1RF pGCP123-$P_{srtA}$-*ftsH*(G37X) | $Fus^r$, $Rif^r$, pGCP123-$P_{srtA}$-*ftsH*(G37X) | This study | BHI with 500 µg/mL kanamycin |
| OG1RF Δ*mprF1* Δ*mprF2* pGCP123-$P_{srtA}$-*ftsH*(G37X) | $Fus^r$, $Rif^r$, Δ*mprF1*Δ*mprF2* pGCP123-$P_{srtA}$-*ftsH*(G37X) | This study | |
| OG1RF pGCP123 | $Fus^r$, $Rif^r$, pGCP123 | This study | |
| OG1RF Δ*mprF1* Δ*mprF2* pGCP123 | $Fus^r$, $Rif^r$, Δ*mprF1*Δ*mprF2* pGCP123 | This study | |
| OG1RF pMSP3535 | $Fus^r$, $Rif^r$, pMSP3535 | This study | BHI with 100 µg/mL erythromycin |
| OG1RF pMSP3535-$P_{nisA}$-6his-*ftsH* | $Fus^r$, $Rif^r$, pMSP3535- $P_{nisA}$-6his-*ftsH* | This study | |
| OG1RF pMSP3535-$P_{nisA}$-2HA-*ftsH* | $Fus^r$, $Rif^r$, pMSP3535-$P_{nisA}$-2HA-*ftsH* | This study | |
| OG1RF pMSP3535-$P_{nisA}$-6his-*ftsH*(H456Y) | $Fus^r$, $Rif^r$, pMSP3535- $P_{nisA}$-6his-*ftsH*(H456Y) | This study | |

| | | | |
|---|---|---|---|
| OG1RF pMSP3535-2HA- P$_{nisA}$-*ftsH*(H456Y) | Fus$^r$, Rif$^r$, pMSP3535- P$_{nisA}$-2HA-*ftsH*(H456Y) | This study | |
| OG1RF pMSP3535-P$_{nisA}$-*cls1* | Fus$^r$, Rif$^r$, pMSP3535- P$_{nisA}$-*cls1* | This study | |
| OG1RF pMSP3535-P$_{nisA}$-RS00500-HA | Fus$^r$, Rif$^r$, pMSP3535- P$_{nisA}$-RS00500-HA | This study | |
| OG1RF pMSP3535-P$_{nisA}$- RS08610-HA | Fus$^r$, Rif$^r$, pMSP3535- P$_{nisA}$-RS08610-HA | This study | |
| OG1RF pMSP3535-P$_{nisA}$- RS02510-HA | Fus$^r$, Rif$^r$, pMSP3535- P$_{nisA}$-RS02510-HA | This study | |
| OG1RF Δ*mprF1*Δ*mprF2 ftsH*(G37X) pMSP3535-P$_{nisA}$-RS00500-HA | Fus$^r$, Rif$^r$, Δ*mprF1*Δ*mprF2 ftsH*(G37X), pMSP3535-P$_{nisA}$-RS00500-HA | This study | |
| OG1RF Δ*mprF1*Δ*mprF2 ftsH*(G37X) pMSP3535-P$_{nisA}$- RS08610-HA | Fus$^r$, Rif$^r$, Δ*mprF1*Δ*mprF2 ftsH*(G37X), pMSP3535- P$_{nisA}$-RS08610-HA | This study | |
| OG1RF Δ*mprF1*Δ*mprF2 ftsH*(G37X) pMSP3535-P$_{nisA}$- RS02510-HA | Fus$^r$, Rif$^r$, Δ*mprF1*Δ*mprF2 ftsH*(G37X), pMSP3535- P$_{nisA}$-RS02510-HA | This study | |
| OG1RF Δ*mprF1*Δ*mprF2* DAP-passage control pMSP3535-P$_{nisA}$-RS00500-HA | Fus$^r$, Rif$^r$, Δ*mprF1*Δ*mprF2,* pMSP3535- P$_{nisA}$-RS00500-HA. passage control strain | This study | BHI with 100 µg/mL erythromycin |
| OG1RF Δ*mprF1*Δ*mprF2* DAP-passage control pMSP3535-P$_{nisA}$- RS08610-HA | Fus$^r$, Rif$^r$, Δ*mprF1*Δ*mprF2,* pMSP3535- P$_{nisA}$-RS08610-HA. passage control strain | This study | |
| OG1RF Δ*mprF1*Δ*mprF2* DAP-passage control pMSP3535-P$_{nisA}$- RS02510-HA | Fus$^r$, Rif$^r$, Δ*mprF1*Δ*mprF2,* pMSP3535- P$_{nisA}$-RS02510-HA. passage control strain | This study | |
| OG1RF RS05275::Tn (*yckE*::Tn) | Fus$^r$, Rif$^r$, RS05275::Tn (Strain ID: 007A07, 114D11, 013G10) | From a transposon library gifted by Gary Dunny, University of Minnesota | BHI |
| OG1RF RS04635::Tn (*lutA*::Tn) | Fus$^r$, Rif$^r$, RS04635::Tn (Strain ID: 030F11, 070D04, 070C04) | | |

| | | | |
|---|---|---|---|
| OG1RF RS07835::Tn (*gelE*::Tn) | Fus$^r$, Rif$^r$, RS07835::Tn (Strain ID: 100E05, 148F04) | (Kristich et al., 2008) | |
| OG1RF RS12410::Tn (*trePP*::Tn) | Fus$^r$, Rif$^r$, RS12410::Tn (Strain ID: 064F10, 095E07, 018D03, 110D12) | | |
| OG1RF RS07350::Tn (*carB*::Tn) | Fus$^r$, Rif$^r$, RS07350::Tn (Strain ID: 006D07, 034D02) | | |
| OG1RF RS07135::Tn (*cryZ*::Tn) | Fus$^r$, Rif$^r$, RS07135::Tn (Strain ID: 040B07) | | |
| OG1RF RS05580::Tn (*hrcA*::Tn) | Fus$^r$, Rif$^r$, RS05580::Tn (Strain ID: 075A04, 016F09) | | |
| OG1RF pMSP3545-*dcas9* pABG5.2mini | Fus$^r$, Rif$^r$, pMSP3535-*dcas9* pABG5.2mini | This study | |
| OG1RF Δ*mprF1*Δ*mprF2* pMSP3545-*dcas9* pABG5.2mini | Fus$^r$, Rif$^r$, Δ*mprF1*Δ*mprF2* pMSP3535-*dcas9* pABG5.2mini | This study | |
| OG1RF pMSP3545-*dcas9* pABG5.2mini-*lysS*_g1 | Fus$^r$, Rif$^r$, pMSP3535-*dcas9* pABG5.2mini-*lysS*_g1 | This study | BHI with 100 µg/mL erythromycin, 500 µg/mL kanamycin |
| OG1RF Δ*mprF1*Δ*mprF2* pMSP3545-*dcas9* pABG5.2mini-*lysS*_g1 | Fus$^r$, Rif$^r$, Δ*mprF1*Δ*mprF2* pMSP3535-*dcas9* pABG5.2mini-*lysS*_g1 | This study | |
| OG1RF pMSP3545-*dcas9* pABG5.2mini-*pyrB*_g1 | Fus$^r$, Rif$^r$, pMSP3535-*dcas9* pABG5.2mini-*pyrB*_g1 | This study | |
| OG1RF Δ*mprF1*Δ*mprF2* pMSP3545-*dcas9* pABG5.2mini-*pyrB*_g1 | Fus$^r$, Rif$^r$, Δ*mprF1*Δ*mprF2* pMSP3535-*dcas9* pABG5.2mini-*pyrB*_g1 | This study | |
| OG1RF pGCP213-*ftsH*(G37X) | Fus$^r$, Rif$^r$, pGCP213-*ftsH*(G37X) | This study | |
| OG1RF Δ*mprF1*Δ*mprF2* pGCP213-*ftsH*(G37X) | Fus$^r$, Rif$^r$, Δ*mprF1*Δ*mprF2* pGCP213-*ftsH*(G37X) | This study | BHI with 50 µg/mL erythromycin |
| OG1RF pGCP213-Δ*dnaK* | Fus$^r$, Rif$^r$, pGCP213-*Δ*dnaK* | This study | |

61

**Table S2. gBlock sequences and PCR primers**

| gBlock | Sequence |
|---|---|
| *lysS_g1* | TATCGACGGAAGATCCTGCAGAGATCTATCTAAAACAGTCTTAATTCTATC<br>TTGAGAAAGTATTGGTAATAATATTATTGTCGATAACGCGAGCATAATAAA<br>CGGCTCTGATTAAATTCTGAAGTTTGTTAGATACAATGATTTCCGATCGAA<br>ACGTTTACCGAAGTTTTAGAGCTAGAAATAGCAAGTTAAAATAAGGCTAGT<br>CCGTTATCAACTTGAAAAAGTGGCACCGAGTCGGTGCTTTTTTTGGATCC<br>ACTAGTGGTACCGAATTCAACCCGAACAATTGGCATGCGGCCGCCACCG<br>CGGT |
| *pyrB_g1* | TATCGACGGAAGATCCTGCAGAGATCTATCTAAAACAGTCTTAATTCTATC<br>TTGAGAAAGTATTGGTAATAATATTATTGTCGATAACGCGAGCATAATAAA<br>CGGCTCTGATTAAATTCTGAAGTTTGTTAGATACAATGATTTCTAACCCCA<br>TGACTTCACGGTGTTTTAGAGCTAGAAATAGCAAGTTAAAATAAGGCTAGT<br>CCGTTATCAACTTGAAAAAGTGGCACCGAGTCGGTGCTTTTTTTGGATCC<br>ACTAGTGGTACCGAATTCAACCCGAACAATTGGCATGCGGCCGCCACCG<br>CGGT |

| Primer | Target(s) | Sequence | Purpose / Remarks |
|---|---|---|---|
| M13F | - pCYW2-*ftsH*(G37X)<br>- pABG5.2mini-*lysS*_g1<br>- pABG5.2mini-*pyrB*_g1<br>- pGCP213-*ftsH*(G37X) | GTAAAACGA<br>CGGCCAG | Screening for transformants targeting the insert<br><br>Same primers used for Sanger sequencing |
| M13R | | CAGGAAACA<br>GCTATGAC | |
| pMSP3535-Screen_R | - pMSP3535-6his-*ftsH*<br>- pMSP3535-2HA-*ftsH*<br>- pMSP3535-6his-*ftsH*(H456Y)<br>- pMSP3535-2HA-*ftsH*(H456Y)<br>- pMSP3535-*cls1*<br>- pMSP3535-RS00500-HA<br>- pMSP3535-RS08610-HA<br>- pMSP3535-RS02510-HA | CGAAATTAAT<br>ACGACTCAC<br>TATAGGG | Screening for transformants targeting the insert<br><br>Same primers used for Sanger sequencing |
| pMSP3535-Screen_F | | TTTTGAAAAC<br>CGCTACGGA<br>TC | |
| pMSP3545-Screen-F | pMSP3545-*dcas9* | TAATACGAC<br>TCACTATAG<br>GG | Screening for transformants targeting the insert<br><br>Same primers used for Sanger sequencing |
| pMSP3545-Screen-R | | GGTTGCAAA<br>TTTTGAAAAC<br>CGC | |
| Amp_FtsH_F | *E. faecalis* colonies from strains obtained from *in vitro* evolution | AGCTGACTA<br>CGTAGGGTT<br>TG | Screening of FtsH locus for truncation of FtsH size on the genome |
| Amp_FtsH_R | | GTGCAGTAT<br>TCGTCAACT<br>CG | |

| | | | |
|---|---|---|---|
| Nil | pGCP213 | Nil | EcoRI and NotI restriction enzymes used to linearise instead |
| pMSP3535-BamHI-F | pMSP3535 | GATCCATGCAGAGTCTCCTGTTTTACAACCGGGTGTACATAGCGAAATACTTGTAATGCGTGGT | To linearize plasmid |
| pMSP3535-PstI-R | | GGAATTCGCATGCGAGCTCGTCGACAGCGCTTCTAGAC | |
| Nil | pCYW2 | Nil | EcoRI and NotI restriction enzymes used to linearise instead |
| 5.2mini_Linearise_F | pABG5.2mini | GGCCGCCACCGCGGTGGAGCTC | To linearize plasmid |
| 5.2mini_Linearise_R | | GATCTTCCGTCGATACTATGTTATACG | |
| FtsH_STOP_Cmpl_F | Genomic DNA of DAP$^R$ strain from *in vitro* evolution: DAP#50 | GCTTGATATCGAATTAATGATGAGCATAAGGAGGA | To create insert: *ftsH*(G37X) |
| FtsH_STOP_Cmpl_R | | ACCGCGGTGGCGGCCTTATTTATAACGATCTTCGTAG | |
| 6his-FtsH_F | *E. faecalis* OG1RF gDNA | GACTCTGCATGGATCATGCATCATCACCATCACCACATGAGCATAAGGAGGACAGG | To create insert: 6his-*ftsH* |
| FtsH_R | | TCGCATGCGAATTCCCTGCACCTTCTACTAATTGGTTATT | |
| 2HA-FtsH_F | *E. faecalis* OG1RF gDNA | GACTCTGCATGGATCATGTACCCATACGATGTTCCAGATTACGCTTACCCATACGATGTTCCAGATTACGCT | To create insert: 2HA-*ftsH* |

| | | | |
|---|---|---|---|
| | | AGCATAAGG AGGACAGGC | |
| FtsH_R | | TCGCATGCG AATTCCCTG CACCTTCTA CTAATTGGTT ATT | |
| (1) 6his-FtsH_F | First PCR template is *E. faecalis* OG1RF gDNA<br><br>Second PCR template is a 1:1 mix of the PCR products from the first reaction: (1+2) and (3+4) | GACTCTGCA TGGATCATG CATCATCAC CATCACCAC ATGAGCATA AGGAGGACA GG | To create insert: 6his-*ftsH*(H456Y)<br><br>Overlap extension PCR. First PCR using primers (1)+(2), (3)+(4). Second PCR using primers (1)+(4) |
| (2) FtsH(H456Y)_R | | GTGTCCCGC TTCGTAGTA AGCCACC | |
| (3) FtsH_(H456Y)_F | | GGTGGCTTA CTACGAAGC GGGACAC | |
| (4) FtsH_R | | TCGCATGCG AATTCCCTG CACCTTCTA CTAATTGGTT ATT | |
| (1) 2HA-FtsH_F | First PCR template is *E. faecalis* OG1RF gDNA<br><br>Second PCR template is a 1:1 mix of the PCR products from the first reaction: (1+2) and (3+4) | GACTCTGCA TGGATCATG TACCCATAC GATGTTCCA GATTACGCT TACCCATAC GATGTTCCA GATTACGCT AGCATAAGG AGGACAGGC | To create insert: 2HA-*ftsH*(H456Y)<br><br>Overlap extension PCR. First PCR using primers (1)+(2), (3)+(4). Second PCR using primers (1)+(4) |
| (2) FtsH(H456Y)_R | | GTGTCCCGC TTCGTAGTA AGCCACC | |
| (3) FtsH_(H456Y)_F | | GGTGGCTTA CTACGAAGC GGGACAC | |
| (4) FtsH_R | | TCGCATGCG AATTCCCTG CACCTTCTA CTAATTGGTT ATT | |
| arcB_F | *E. faecalis* OG1RF gDNA | GACTCTGCA TGGATCTAG GAGGAATCA TCATGAATTC AGT | To create insert: RS00500-HA |

| | | | |
|---|---|---|---|
| arcB_HA_R | | TCGCATGCGAATTCCCTACTAAGCGTAATCTGGAACATCGTATGGGTACACACGAGGAATGAATAAGTTGC | |
| RS08610_F | *E. faecalis* OG1RF gDNA | GACTCTGCATGGATCTGGAGGAATCAACGAATGAAAAAATT | To create insert: RS08610-HA |
| RS08610_HA_R | | TCGCATGCGAATTCCTTATTAAGCGTAATCTGGAACATCGTATGGGTATTTACTCATTAAGCCATCATGGATTTT | |
| RS02510_F | *E. faecalis* OG1RF gDNA | GACTCTGCATGGATCTGGTATAGGAGGATAAAAATGTCTAAATTTTTA | To create insert: RS02510-HA |
| RS02510_HA_R | | TCGCATGCGAATTCCTTATTAAGCGTAATCTGGAACATCGTATGGGTACTGCTCATCTCTATTTATTTTTTTACTGTTTG | |
| (1) dnaK_up_F | *E. faecalis* OG1RF gDNA | GATATCTGCAGAATTTATGAAGTATCAGGACATGGCAA | To create insert: Δ*dnaK*<br><br>Overlap extension PCR. First PCR using primers (1)+(2), (3)+(4). Second PCR using primers (1)+(4) |
| (2) dnaK_up_R | | CCAGTCCCTAAAATCAATTGTTATTGAAGTGAATATCTCCAATCTGTATTAGTTTT | |
| (3) dnaK_down_F | | AAAACTAATACAGATTGGAGATATTCACTTCAATAACAATTGATTTTAGGGACTGG | |
| (4) dnaK_down_R | | CAGTGTGCTGGAATTTCTGGGTGTGTGCCTG | |

| | | | |
|---|---|---|---|
| dnaK_F | *E. faecalis* colonies obtained after homologous recombination | ACTCTAGAGGATCCCATGAGTAAAATTATTGGTATTGACTTAGGA | Screening Δ*dnaK* colonies to confirm for loss of *dnaK* |
| dnaK_R | | ATTCGAGCTCGGTACTTTGTCATCACCATTTACTTCTTCAAAA | |

62
