## Supplementary text for "An essential protease, FtsH, influences daptomycin resistance acquisition in *Enterococcus faecalis*"

### Supplementary methods

#### Bacterial strains and culture conditions

The wild-type (WT) parental strain used was *E. faecalis* OG1RF (Dunny et al., 1978) and Δ*mprF1*, Δ*mprF2,* Δ*mprF1* Δ*mprF2* mutants used were described previously (Kandaswamy et al., 2013, Chong et al., 2017, Rashid et al., 2023). Unless stated, all bacterial strains were grown overnight to late stationary phase for 16-18 hrs in their respective media at 37 °C in static conditions and stored in 25 % glycerol solution at -80 °C when long-term storage was required. If mid-log phase cultures were required, late stationary phase cultures were sub-cultured 1:10 dilution into fresh media and grown at 37 °C, 250 rpm shaking conditions until an $OD_{600}$ of 0.5 ± 0.05 was reached. When normalization of cultures was required, they were centrifuged at 6,000 x g for 5 mins at 4 °C and cell pellets were washed twice with 1 mL of phosphate buffered saline (PBS). Cell suspensions were then normalized to an $OD_{600}$ of 0.7 or required optical density by diluting with PBS. Strains, their respective genotype, source, and growth media used are listed in **supplementary table S1**.

#### Growth curves

Overnight stationary phase cultures were normalized to $OD_{600}$ of 0.7 and diluted 200-fold before inoculating in 200 µL of media in 96-well plates in a ratio of 1:25. The 96-well plates were incubated at 37 °C in a Tecan Infinite© M200 Pro spectrophotometer (Tecan, Switzerland) with absorbance read at 600 nm every 15 mins for 18 hrs.

#### Live/dead staining

Late stationary and mid-log phase cultures were normalized to $OD_{600}$ 0.5 in 1 mL of PBS and washed twice with PBS. 2 µL of SYTO9 and propidium iodide (PI) mix (LIVE/DEAD™ BacLight™ Bacterial Viability Kit, for microscopy, Invitrogen, USA) were added to the cell suspensions and incubated at 15 mins at room temperature in the dark. Stained cells were washed once with PBS and resuspended in 200 µL 0.01 M phosphate buffer (PB). 5 µL of stained cells were wet mounted on 1.0-1.2 mm microscope slides (Biomedia, Singapore) with 0.13-0.16 mm coverslips. Slides were imaged with a Zeiss Axio observer Z1 inverted microscope (Carl-Zeiss, Germany) with a 100X oil immersion objective (NA 1.4, optovar 1.6X), AF488/FITC filter cube (460-490 nm band pass excitation filter, 515-550nm band pass barrier filter) and AF568/Cy3 filter cube (530-550nm band pass excitation filter, 590nm long pass barrier filter). Phase contrast, green (460-490 nm) and red (530-550nm) fluorescence images were captured.

**RNA isolation and sequencing**

Overnight cultures were subcultured 1:10 in BHI and grown to mid-log phase, cultures were then induced for expression of their respective plasmids' gene constructs with 125 ng mL$^{-1}$ of nisin for 16-18 hrs at 37 °C in static conditions. 2 mL of RNAprotect© Bacteria Reagent (Qiagen, Germany) was added to 1 mL of induced culture and incubated for 5 mins at room temperature. Cells were then pelleted and rinsed with PBS to thoroughly remove the RNAprotect© reagent. Cell pellets were then resuspended in 20 mg mL$^{-1}$ lysozyme (Sigma-Aldrich, USA) in lysis buffer (10 mM Tris-HCl pH 7.0, 1 mM EDTA, 50 mM NaCl and 0.74 M Sucrose) and incubated for 1 hr at 37 °C. Cells were then pelleted and washed once with PBS. 1 mL of ice-cold TRIzol™ Reagent (Ambion, USA) was next added and the cell pellet resuspended by pipetting. 200 µL of ice-cold chloroform (Fisher, USA) was next added to the tube and the suspension mixed by shaking gently before incubating on ice for 2 mins. The mixture was then centrifuged for 15 mins at 12, 000 x g at 4 °C to induce phase separation. Next, 500 µL of the upper aqueous phase was carefully removed, mixed with 500 µL of ethanol and added to the spin-column from the PureLink™ RNA Mini Kit (Thermofisher, USA). Purification was then carried as specified by the manufacturer's instructions and eluted in 30 µL of Nuclease-free water (Ambion, USA). Quantification of RNA and DNA were performed using Qubit™ RNA Assay Kits and Qubit™ dsDNA HS Assay Kits (Invitrogen, USA), respectively. Integrity of RNA was analyzed by gel electrophoresis using Agilent RNA ScreenTape (Agilent Technologies, USA). Extracted RNA samples were subjected to ribosomal depletion with Ribo-Zero™ Magnetic Kits (Lucigen, USA) and purified using RNAclean® XP beads (Beckman Coulter, USA) according to their respective manufacturers' protocol. cDNA synthesis was done using NEBNext® RNA First Strand Synthesis Module and NEBNext® Ultra Directional RNA Second Strand Synthesis Module (New England Biolabs, USA), and purified using AMPure XP beads (Beckman Coulter, USA). RNA library preparation and sequencing were done by the sequencing facility of Singapore Centre of Life Science Engineering (SCELSE, Singapore) using MiSeq.

Sequencing reads were mapped to the *E. faecalis* OG1RF reference genome (NCBI accession: NC_017316.1) using BWA (version 0.5.9) on default settings (Li and Durbin, 2009, Nagalakshmi et al., 2010). Reads mapping onto predicted open reading frames (ORFs) were counted on HTseq and ribosomal reads were filtered out (Anders et al., 2015). Pairwise comparisons were performed in in R program with the Bioconductor package, edgeR (Robinson et al., 2010). Significant differentially expressed genes were determined using a cutoff of false discovery rate (FDR) and p-value of 0.05. Kyoto Encyclopedia of Genes and Genomes (KEGG) annotation along with manual curation using BLASTP and cross-referencing of other functional data from literature were performed. Gene ontologies were obtained from KEGG (identifier: efi) and genes were classified based on membership of a pathway with their respective log fold change (logFC).

**FtsH substrate verification by western blot**

Cultures were grown to mid-log phase in BHI (OD$^{600}$ 0.500 ± 0.050) and spiked with 400 µg mL$^{-1}$ of spectinomycin. Cultures were incubated at 37 °C, 200 rpm in shaking conditions where 1 mL of culture was collected at different time points. Culture aliquots were then pelleted and lysed in 20 mg mL$^{-1}$ lysozyme in lysis buffer (10 mM Tris-HCl pH 7.0, 1 mM EDTA, 50 mM NaCl and 0.74 M Sucrose) supplemented with 400 µg mL$^{-1}$ of spectinomycin for 30 minutes at 37 °C. 10 µL of 1M DTT and 33.34 µL of 4X LDS sample loading buffer were added and boiled on a heat block at 100 °C for 15 mins, or until the solution becomes clear. SDS-PAGE and western blot were performed on these samples as described in a previous study (Nielsen et al., 2012). 4-12 % NuPAGE® Bis-Tris mini gel in a XCell SureLock® Mini-Cell filled with either 1X MES or 1X MOPs SDS running buffer (Invitrogen, USA) were used and ran at 140 V for 90 mins. Proteins were transferred to nitrocellulose membranes using the iBlot™ Dry Blotting System (Invitrogen, USA) according to the manufacturer's protocol. Mouse anti-HA (Thermoscientific, USA) was used as the primary antibody at 1:1000 dilution, and goat anti-mouse HRP (Thermoscientific, USA) was used as the secondary antibody at 1:5000 dilution. SuperSignal™ West Femto Maximum Sensitivity Substrate was used as the developing solution at luminol:peroxide:water (1:1:8) ratio.

**ATP quantification assay**

Overnight cultures were normalized to OD$_{600}$ 0.5 in PBS. 100 µL of normalized cultures were then added to wells of an opaque walled 96-well microtiter plate with 100 µL of BacTiter-Glo™. The plate was then incubated for 5 minutes on a shaker at room temperature and luminescence was measured using a Tecan Microplate reader.

**Molecular cloning**

The respective plasmids were extracted from *E. coli* strains harboring them using the Monarch® Plasmid Miniprep Kit (New England BioLabs, USA) according to the manufacturer's instructions.

To generate constructs and ligate inserts into vectors, plasmids were first linearized by PCR and inserts generated by PCR using *E. faecalis* OG1RF genomic DNA as a template. PCR was done using Q5® High-Fidelity DNA Polymerase (New England BioLabs, USA) according to the manufacturer's instructions. Linearized plasmids and their corresponding inserts were then ligated using the In-Fusion HD Cloning system (Takara, Japan) and transformed into Stellar™ Competent Cells (Takara, Japan) according to the manufacturer's instructions.

Plasmids were then extracted from the stellar competent cells using the Monarch® Plasmid Miniprep Kit and then transformed into electrocompetent *E. faecalis*. PCR to create *ftsH*(H456Y) and Δ*dnaK* related inserts were done by overlap extension PCR. To encourage and select for homologous recombinants, transformants of the pGCP213 temperature sensitive vector were grown in the presence of erythromycin at 30 °C to select for chromosomal integrants. To encourage plasmid excision by homologous recombination, selected integrants where than serial passaged at 37 °C in the absence of erythromycin. Erythromycin sensitive colonies were then subjected to PCR to detect the loss of the target gene.

CRISPRi knockdown strains were created as previously described (Afonina et al., 2020). pABG5.2mini plasmid was first linearized by PCR and ligated with inserts containing the guide RNA by using the In-Fusion HD Cloning system. These inserts were in the form of gBlocks ordered from Integrated DNA Technologies Pte. Ltd, Singapore. gBlock (insert) sequences can be found in the **supplementary table S2**. pMSP3545-dcas9 was first transformed into *E. faecalis* wild type and ∆*mprF1* ∆*mprF2*, followed by the respective pABG5.2mini based guide RNA plasmids.

Transformants were selected by BHI or LB agar containing the appropriate antibiotics as listed in **supplementary table S1**. All screening of transformants were done using colony PCR using Taq DNA Polymerase, recombinant (5 U μL$^{-1}$) (Thermo Scientific, USA) according to the manufacturer's instructions. Gel electrophoresis to assess product sizes were done using 1% w/v agarose gel in TAE buffer ran at 100 V for 30 mins followed by ethidium bromide staining for 10-15 mins. After each transformation step, if plasmids passed the colony PCR check, they were extracted and sent for Sanger sequencing for their inserts to ensure the correct sequence is present (1st BASE DNA Sequencing Services, Singapore). PCR purification was done using Wizard® SV Gel and PCR Clean-Up System (Promega, USA) according to the manufacturer's instructions. Sequences of the primers used for PCR are shown in **supplementary table S2**.
